# Global transcriptome analysis identifies nicotinamide metabolism to play key roles in IFN-γ and nitric oxide modulated responses

**DOI:** 10.1101/2025.01.17.633516

**Authors:** Avik Chattopadhyay, Sai Shyam, Shreyasee Das, Dipankar Nandi

## Abstract

Interferon-gamma (IFN-γ) is a key regulator of immune responses. A hallmark of the IFN-γ response is inducible nitric oxide (NO) production, driven primarily by nitric oxide synthase 2 (NOS2). In this study, we investigated the influence of NO on the IFN-γ-induced transcriptomic and metabolic changes in the RAW 264.7 macrophage cell line. IFN-γ activation led to NO-dependent lactate production and lower cell survival. Bulk RNA sequencing analysis identified genes differentially expressed early by IFN-γ that were either NO-independent or NO-dependent. Inhibition of NO modulated a minor subset of the transcriptome, notably affecting *Klf6* (a tumor suppressor) and *Zfp36* (a regulator of pro-inflammatory cytokines). Interestingly, both *Klf6* and *Zfp36* correlatively showed high expression in most cancers. The PPI network exhibited dense clustering with scale-free and small-world properties, identifying *Stat1*, *Irf7*, *Irf1*, *Cxcl10*, and *Isg15* as top five hubs. The top IFN-γ signalling genes exhibited deficient expression in the brain but were highly expressed in lung, spleen, and EBV-transformed lymphocytes. Gene-disease associations linked the IFN-γ-regulated genes to immunodeficiencies, chronic inflammatory responses and malignancies. Interestingly, IFN-γ upregulated genes were involved in nicotinamide metabolism, suggesting a transcriptional basis for modulation of metabolic pathways. This novel aspect was experimentally tested to show that IFN-γ induced NAD amounts. Importantly, the inhibition of purine nucleoside phosphorylase (using Forodesine hydrochloride) which is involved in the endogenous pathway for NAD generation, lowered IFN-γ induced nitrite and increased cell survival *in vitro*. Functionally, enriched nicotinamide metabolism by IFN-γ may regulate inflammatory responses and the implications of our findings are discussed.

## Introduction

Interferon-gamma (IFN-γ), a type II interferon, is a pivotal cytokine in orchestrating immune responses, particularly in host defense against pathogens and regulating inflammatory processes. IFN-γ acts as a macrophage activating factor by polarizing macrophages to M1 state, stimulating these cells to become more effective in eliminating pathogens and promoting immune responses against cancer. Macrophages undergo profound transcriptomic changes upon IFN-γ stimulation, driving shifts in cellular metabolism, effector functions, and immune responses [1]. Therefore, the intricate network of molecular events triggered by IFN-γ activation in macrophages is a subject of intense investigation, including its transcriptional regulation.

Nitric Oxide Synthases (NOS) are enzymes that produce nitric oxide (NO) from L-arginine and oxygen, requiring multiple cofactors such as tetrahydrobiopterin (BH4), Nicotinamide adenine dinucleotide phosphate (NADPH), Flavin adenine dinucleotide (FAD), flavin mononucleotide (FMN), and heme. Nitric Oxide Synthase 2 (NOS2), also known as inducible NOS (iNOS), is an isoform activated by IFN-γ, operates independently of calcium unlike NOS1 and NOS3, and generates higher NO levels. IFN-γ-induced NOS2 function plays a crucial role in macrophage activation by significantly increasing NO production [2].

Previously, we demonstrated that IFN-γ stimulates NOS2-mediated NO production, leading to increased ROS levels in mouse cell lines: H6 hepatoma and L929 fibrosarcoma. This IFN-γ-induced NO and ROS production triggers activation of the JNK signaling pathway, culminating in cell cycle arrest and apoptosis [3, 4]. Our recent publication using metabolic flux analyses revealed that IFN-γ-activated H6 cells undergo metabolic reprogramming, suppressing mitochondrial oxygen consumption while augmenting glycolytic flux, resulting in elevated glucose uptake and lactate production. Notably, IFN-γ-induced NO and ROS production and enhanced glycolysis and lactate production exhibit reciprocal regulation, collectively influencing tumor cell growth [5]. Similarly, IFN-γ-activated RAW 264.7 macrophages demonstrate heightened nitric oxide (NO) and reactive oxygen species (ROS) production, concurrent augmentation in glycolytic activity, and elevated lactate production [6, 5].

NO, via its engagement with heme and non-heme metal-containing proteins, alongside constituents of the electron transport chain, serves as a dual agent: regulating cellular respiration and modulating intracellular metabolism. Furthermore, the multifaceted impacts of NO and its derivatives, reactive nitrogen species (RNS), entail intricate interactions with various targets contingent upon concentration, as well as temporal and spatial constraints [7]. Also, NO facilitates the aggregation of activated macrophages by upregulating Cd11b expression, critical for motility and phagocytic functions and orchestrating a coordinated defense against pathogens [8]. It eliminates intracellular pathogens such as *Mycobacterium* and *Salmonella* through interfering with bacterial respiration, directly damaging bacterial DNA, proteins, and lipids, leading to pathogen death [9]. Additionally, NO triggers apoptosis in host cells harboring pathogens, aiding in the clearance of infected cells [10]. NO protects against bacterial sepsis, as evidenced by observations as *Nos2*^-/-^ mice display diminished survival in the face of bacterial sepsis [11, 12]. Furthermore, NO exerts significant antiviral effects, notably impeding the proliferation of pathogens such as SARS-CoV-2, safeguarding cellular integrity against viral-induced cytotoxicity and protecting patients in COVID-19 pneumonia through accelerated nasal viral clearance [13, 14, 15].

However, the dual nature of NO also contributes to inflammatory and autoimmune diseases, presenting both a defender and a provocateur. Pharmacological interventions targeting IFN-γ-induced NO production ameliorates conditions like ulcerative colitis and sepsis [16]. These findings underscore the impact of IFN-γ-induced NOS2-dependent NO production on various cellular processes, including glycolytic enhancement and cellular growth, determining health and disease outcomes. However, limited knowledge is available on the impact of NOS2-mediated NO on IFN-γ-induced transcriptional processes, its role in metabolic reprogramming, and the functional dynamics of the IFN-γ signaling network from a graph theory perspective. In this investigation, we aimed to elucidate IFN-γ-induced NO modulation of macrophage transcriptome in response to IFN-γ stimulation. Our methodology involved the stimulation of RAW 264.7 macrophages with IFN-γ, concomitantly administering N^ω^-methyl L-arginine (LNMA), an inhibitor targeting all NOS isoforms, thus impeding NO production. Leveraging RNA sequencing (RNA-seq) technology, we aimed to unravel the intricate interplay between IFN-γ signaling and NOS-mediated modulation of transcript levels post 6 hr of induction. In addition, we unraveled the intricacies of the IFN-γ signaling-induced transcriptome by focusing on its protein-protein interaction (PPI) network. In addition, we compared gene expression across tissues and identified diseases associated with the top-upregulated genes to delineate potential therapeutic targets. Interestingly, we identified several genes involved in nicotinamide metabolism to be IFN-γ-inducible and experimentally tested the functional roles of NAD in our model system. The implications of this novel observation with regard to host inflammatory responses are discussed. Overall, our study sought to comprehensively characterize IFN-γ signaling in macrophages, elucidating its impact on cellular functions, and exploring its associations with different diseases.

## Results

### IFN-γ activated RAW 264.7 macrophages produce elevated amounts of lactate in NO-dependent manner

Our study investigated the roles of nitric oxide (NO) in the glycolytic metabolic reprogramming of macrophages activated by IFN-γ. We used RAW 264.7 macrophages treated with IFN-γ and inhibited nitric oxide synthases (NOS) using LNMA (a pan-NOS inhibitor) and 1400W (a NOS2-specific inhibitor). Both inhibitors reduced nitrite levels (an indicator of NO production) and rescued cell numbers, confirming effective NOS inhibition. IFN-γ activation significantly increased lactate release, indicating enhanced glycolysis. Inhibition of NOS resulted in a partial but significant reduction in lactate levels (Fig. 1). Overall, these observations demonstrated that the elevation of lactate production to IFN-γ response is NO-dependent.

**Figure 1.**
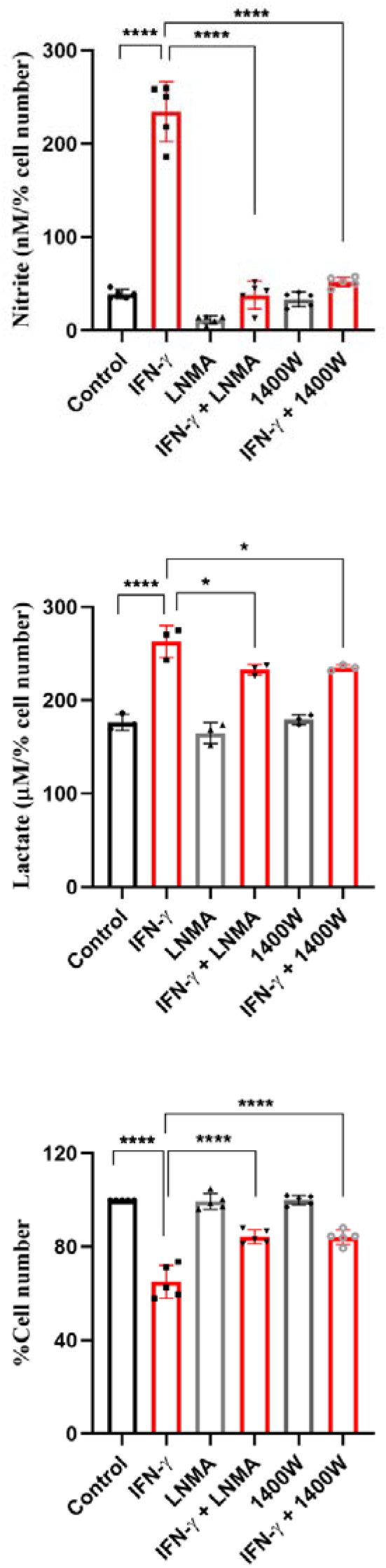
IFN-γ activated RAW 264.7 macrophages produce enhanced levels of lactate, partially in NO-dependent manner. The concentration of nitrite and lactate in cell-free supernatant was estimated using colorimetric assays. The percentage of cell number was measured using Trypan blue dye exclusion assay. All data were acquired after 24 hours of activation. Statistical analyses were performed using ordinary one-way ANOVA with Sidak’s multiple comparisons tests. (*) and (****) indicate the statistical differences of p < 0.05 and p < 0.0001 between the compared groups. Each data point is representative of an independent experiment. Data are represented as mean ± SD of 3-5 independent experiments.

### Nitric Oxide has a minor influence on Interferon-gamma-induced differential gene expression, notably affecting *Klf6* and *Zfp36*

Given the critical role of NO in IFN-γ-induced enhancement of glycolytic metabolism (5), we further explored its impact on shaping the macrophage transcriptome. We conducted bulk RNA sequencing on RAW 264.7 macrophages across three independent experiments under four conditions: untreated, IFN-γ treated, LNMA treated, and IFN-γ plus LNMA treated. This analysis aimed to elucidate the early transcriptomic changes mediated by NO in IFN-γ-activated macrophages. A peak at high Phred scores indicated most high-quality reads, suggesting good sequencing data with accurate base calls. High Phred scores at each position along the reads indicate accurate base calls. Therefore, the RNA-seq data is reliable for downstream analysis (Fig. S1). The QC passed reads were mapped onto indexed Mouse reference genome (GRCm 38.90) using STAR v2 aligner. On average 92.76% of the reads aligned onto the reference genome (Supplementary Tables 1 and 2). For the scope of this study, we focused on the differential expression of only protein-coding genes. Among downregulated genes, exclusively, 27 genes were for IFN-γ and 14 genes for IFN-γ with LNMA. Among upregulated genes, exclusively, 93 genes for IFN-γ-activation, 131 genes for LNMA treatment, and 45 genes for the combination of IFN-γ and LNMA. 34 genes were shared between IFN-γ activation and the combination of IFN-γ and LNMA. The top 10 IFN-γ-induced upregulated protein-coding genes were *Gm17334*, *Irgm1*, *Acod1*, *Cxcl10*, *Irf1*, *Isg15*, *H2-T24*, *Ifitm3*, *Hcar2* and *Mpeg1*. The top 5 IFN-γ-induced downregulated protein-coding genes were *4930516K23Rik*, *Fam26d*, *Rhox2g*, *Olfr173*, and *Lypd2*. The top 5 LNMA-induced upregulated protein-coding genes were *Gm14444*, *Tmem37*, *Hist1h4k*, *Tas1r1*, and *Tpsab1*. The top 5 LNMA-induced downregulated protein-coding genes were *Saa2*, *Msmp*, *B130006D01Rik*, *Zfp41*, and *Dhrs11*. The top 5 IFN-γ and LNMA-combination-induced upregulated protein-coding genes were *Isg15*, *AC125444.5*, *Psme2*, *AC157931.1*, and *Psme1*. The top 5 IFN-γ and LNMA-combination-induced downregulated protein-coding genes were *Cd300ld2*, *Tspo2*, *Hmox1*, *Esco2*, and *Gm8127* (Fig. S2).

An analysis of NOS-regulated differential transcript levels revealed that inhibiting NOS using LNMA in the presence of IFN-γ signaling did not influence the transcript level of any remarkable cluster of genes (Fig. 2A, S3, S4). The DEGs were categorized into NOS-independent and NOS-dependent groups, based on the effect on transcript levels with or without LNMA treatment upon IFN-γ activation. Our analysis revealed that 34 genes were upregulated and 1 gene was downregulated upon IFN-γ activation in NOS-independent manner. 4 genes (*Klf6*, *Zfp36*, *Inha*, and *Rbm43*) were upregulated and 6 genes (*Scgb2b2*, *Tcp10c*, *Vip*, *Il11ra2*, *Hsd17b3*, and *Fndc9*) were downregulated upon IFN-γ activation, which were not significantly affected upon IFN-γ and LNMA combination treatment, marking them genes whose transcript levels were differentially regulated upon IFN-γ treatment in NOS-dependent manner (Table 1).

**Figure 2.**
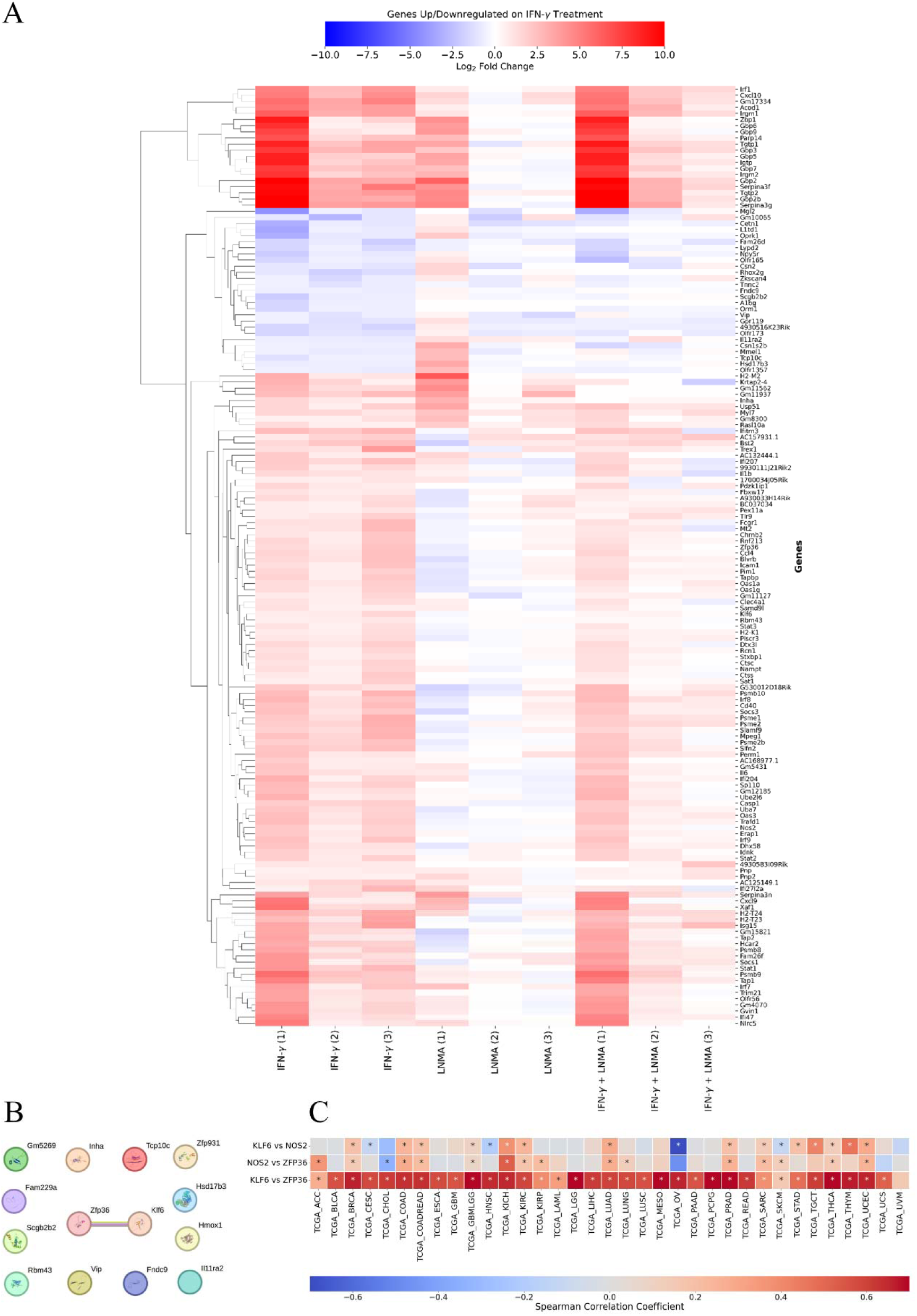
Inhibition of NO during early IFN-γ activation affects a small subset of genes, notably *Klf6* and *Zfp36*. (A) Heatmap displaying the log2(fold change) values with respect to control for the IFN-γ-induced DEGs. The hierarchical clustering was performed based on IFN-γ-induced DEGs. Data is representative of three independent experiments. (B) PPI network of IFN-γ-induced NO-dependent genes as constructed on String of EMBL-EBI at a confidence score cut-off of 0.4. (C) Heatmap displaying correlation of gene expression between TCGA datasets. * indicates p value ≤ 0.05.

**Table 1.** List of NOS-dependent protein-coding DEGs upon IFN-γ activation and inhibition of NO. The Ensembl IDs have a prefix as ‘ENSMUSG000000.’ The log2 (fold change) values are listed upto two digits after decimal. Combination refers to the combined treatment of IFN-γ and LNMA. The numbers after the comparison group indicate the serial numbers of independent experiments.

| Ensembl ID | Gene name | Control v/s IFN-γ-1 | Control v/s IFN-γ-2 | Control v/s IFN-γ-3 | Mean of Control v/s IFN-γ | Control vs combination-1 | Control vs combination-2 | Control vs combination-3 | Mean of Control vs combination |
| --- | --- | --- | --- | --- | --- | --- | --- | --- | --- |
| 44786 | *Zfp36* | 1.47 | 0.91 | 2.40 | 1.59 | -0.38 | 0.25 | -0.15 | -0.09 |
| 00078 | *Klf6* | 0.75 | 0.81 | 1.60 | 1.05 | 0.39 | 0.47 | 0.11 | 0.32 |
| 32968 | *Inha* | 1.58 | 0.90 | 0.58 | 1.02 | 0 | 0.34 | 0.35 | 0.23 |
| 36249 | *Rbm43* | 0.66 | 0.70 | 0.93 | 0.76 | 0.30 | 0.28 | -0.12 | 0.15 |
| 78554 | *Fam229a* | -2 | 0.52 | 0.27 | -0.39 | 0.58 | 0.87 | 0.88 | 0.77 |
| 36521 | *Scgb2b2* | -2 | -0.77 | -0.62 | -1.13 | -0.41 | 0.18 | 0.13 | -0.03 |
| 52469 | *Tcp10c* | -1.22 | -0.62 | -0.59 | -0.81 | 0.19 | -0.31 | -0.07 | -0.06 |
| 19772 | *Vip* | -0.58 | -1.09 | -0.67 | -0.78 | 0.41 | -0.26 | -0.48 | -0.11 |
| 33122 | *Hsd17b3* | -0.73 | -0.67 | -0.63 | -0.68 | -0.32 | -0.13 | 0.21 | -0.08 |
| 48721 | *Fndc9* | -0.67 | -0.58 | -0.73 | -0.66 | 0.08 | 0 | -0.16 | -0.02 |
| 05413 | *Hmox1* | 0.40 | -0.50 | 1.59 | 0.49 | -0.67 | -0.73 | -1.34 | -0.91 |
| 78861 | *Zfp931* | -0.19 | -0.13 | 0.96 | 0.21 | -0.59 | -0.74 | -0.79 | -0.71 |
| 98549 | *Gm5269* | -0.46 | -0.55 | 0 | -0.34 | -1.58 | -0.77 | -2.70 | -1.68 |

A PPI network construction using the NOS-dependent DEGs revealed that only *Klf6* and *Zfp36* formed an association (Fig. 2B). Assuming that such an association can be functional, we looked for the correlation of gene expression with respect to NOS2 in human cancers of TCGA database. No strong correlation of expression was reported in any cancer type. However, *KLF6* and *ZFP36* showed a strong expression pattern in most cancer types, notably, top five correlation was observed in thyroid cancer, mesothelioma, pheochromocytoma and paraganglioma, prostate adenocarcinoma, and glioma (Fig. 2C).

### Functional enrichment analysis reveals typical features of IFN-γ signaling

For the gene ontology biological process, the smallest Term p-value of 0.00000271 was found for ISG15-protein conjugation, with the genes associated as *Isg15*, *Uba7*, and *Ube2l6*. The second most diminutive term p-value of 0.0000645 was found with Tap-dependent antigen processing and presentation of endogenous peptide antigen via MHC class I via the ER pathway, involving *Tap* (Fig. 3, Fig. S5). Other significant biological processes found were nicotinamide riboside metabolism (*Pnp*, *Pnp2*), ABC-type peptide antigen transporter activity (*Tap1*, *Tap2*), regulation of MyD88-dependent toll-like receptor signaling pathway (*Irf1*, *Irf7*), negative regulation of primary miRNA processing (*Il6*, *Stat3*), negative regulation of dendritic cell cytokine production (*Bst2*), positive regulation of interleukin-21 production (*Il6*), etc. Among cellular components GO terms, the top four significant terms were symbiont-containing vacuole (*Gbp2*, *Gbp2b*, *Gbp3*, *Gbp6*, *Gbp7*), ISGF3 complex (*Irf9*, *Stat1*, *Stat2*), proteasome activator complex (*Psme1*, *Psme2*), and TAP complex (*Tap1*, *Tap2*). The most significant molecular function was ABC-type peptide antigen transporter activity (*Tap1*, *Tap2*). Among immune functions, the three significant terms were toll-like receptor signaling pathway (*Irf1*, *Irf7*), negative regulation of CD8-positive, alpha-beta T cell differentiation and negative regulator of cytokine signalling (*Socs1*), and negative regulation of plasmacytoid dendritic cell cytokine production (*Bst2*) (Fig. 3, Fig. S5).

**Figure 3.**
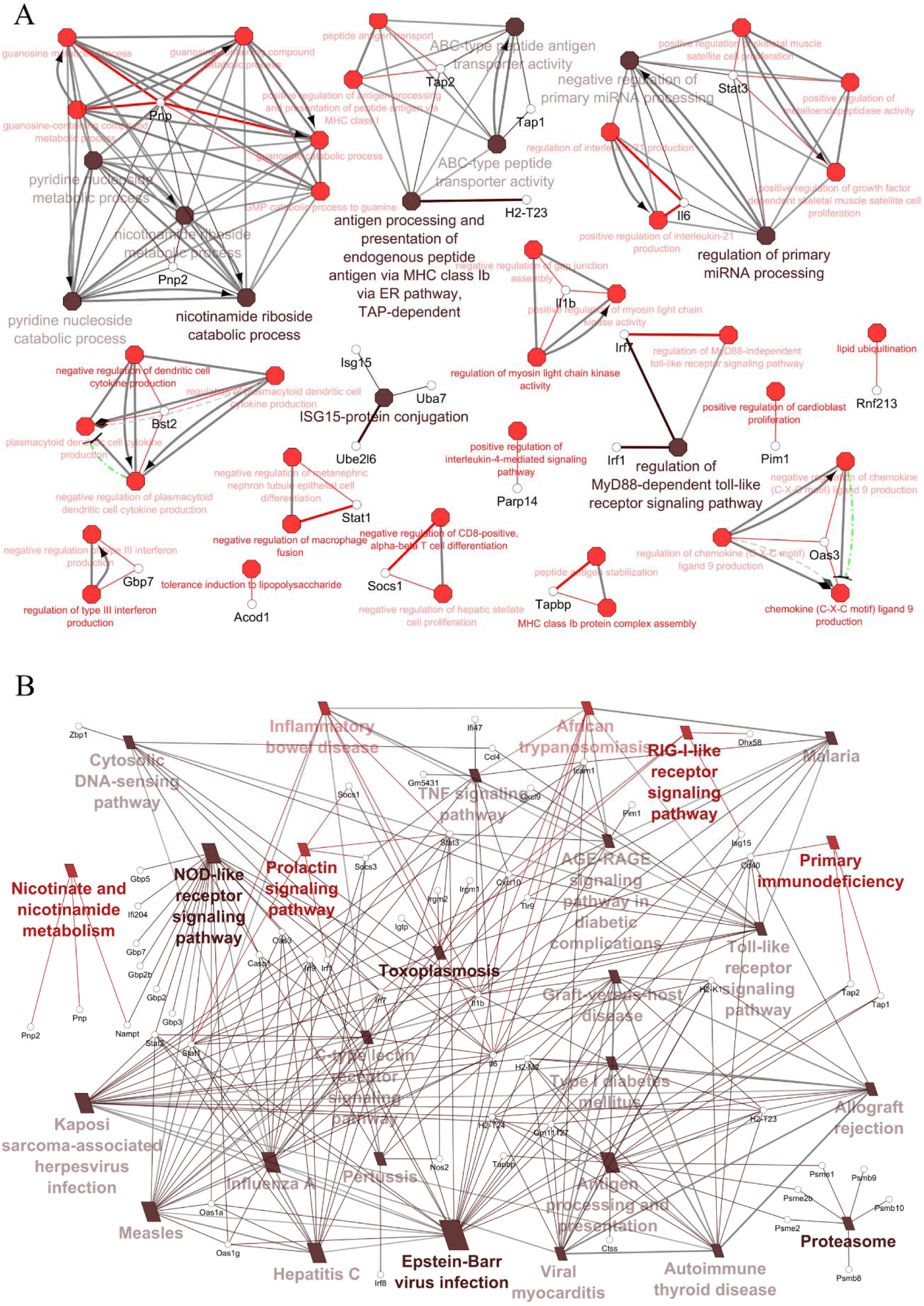
Functional enrichment analysis uncovers the typical features of IFN-γ signaling, especially nicotinamide metabolism. IFN-γ-induced upregulated DEGs were used as input in the ClueGO plugin of Cytoscape. (A) Biological Processes of Gene Ontology and (B) KEGG pathways are displayed. Brown and red color of the nodes represent the Benjamini-Hochberg False Discovery Rate ≤ 0.0001 and ≤0.05, respectively.

### The PPI network of IFN-γ signaling form a scale-free network with small world properties

The protein-protein interaction (PPI) network of IFN-γ-induced upregulated genes was constructed at the highest confidence of 0.9 for minimum required interaction score, to build a very robust network. The network formed with 78 nodes, 275 edges and a clustering coefficient of 0.61. Such a high clustering coefficient indicates that the IFN-γ signaling PPI network form local neighborhoods of densely interconnected nodes (Fig. 4A). The topological properties of the network are tabulated in Table 2. The degree, betweenness centrality and log_2_(fold change) were compared with respect to each other. Log2(fold change) exhibited no correlation with degree and betweenness centrality, indicating, nodes with the greatest changes in expression between groups may not have the biggest influence on other nodes within the network. On the other hand, degree and betweenness centrality showed significant correlation (Spearman rank correlation coefficient = 0.89, p-value < 0.0001) (Fig. 4B).

**Figure 4.**
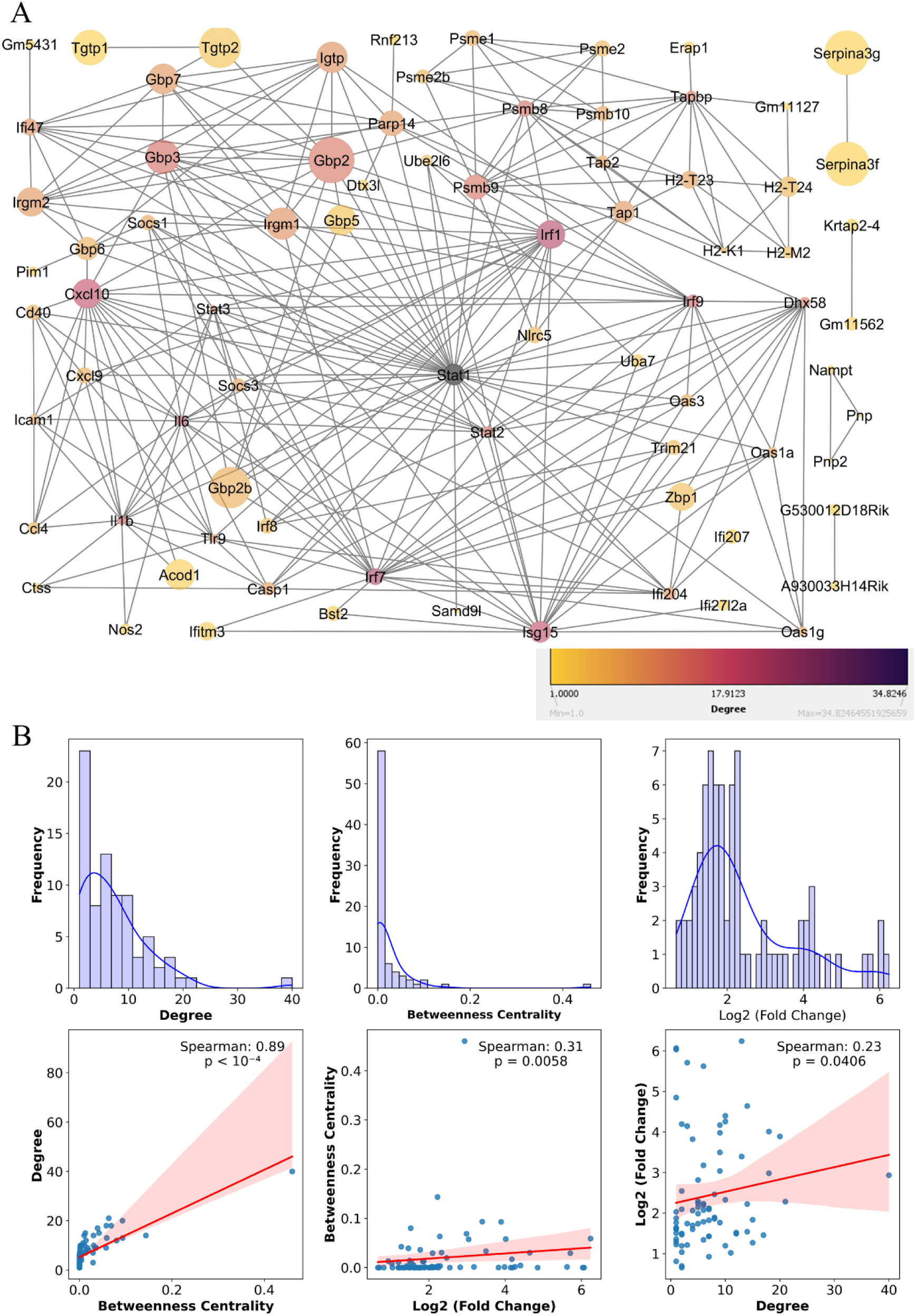
The PPI network of IFN-γ signaling displays robust clustering with a significant positive correlation between node degree and betweenness centrality. (A) The PPI network was constructed using the significantly upregulated DEGs of IFN-γ signaling. The gene names were imported in the STRING tool of EMBL-EBI and the PPI network was created with the highest confidence cut-off of 0.9. The node sizes are proportional to the fold change values in RNA-seq. The node colors represent the degree values. (B) The KDE plot demonstrates the relationship between degree, betweenness centrality of the PPI network and log2 (fold change) values of the nodes from RNA-seq data.

**Table 2.** Expression and topological properties of hubs in the IFN-γ signaling PPI network.

| Name | Degree | Betweenness centrality | Mean log<sub>2</sub> (fold change) | Mean shortest pathlength | Closeness centrality | Clustering coefficient | Topological coefficient |
| --- | --- | --- | --- | --- | --- | --- | --- |
| *Stat1* | 40 | 0.46 | 2.93 | 1.49 | 0.67 | 0.18 | 0.15 |
| *Irf7* | 21 | 0.06 | 2.28 | 2.01 | 0.50 | 0.30 | 0.22 |
| *Irf1* | 20 | 0.09 | 3.89 | 1.84 | 0.54 | 0.33 | 0.22 |
| *Cxcl10* | 18 | 0.06 | 4.01 | 2.03 | 0.49 | 0.37 | 0.25 |
| *Isg15* | 18 | 0.07 | 2.99 | 2.09 | 0.48 | 0.32 | 0.24 |
| *Il6* | 17 | 0.04 | 1.46 | 2.07 | 0.48 | 0.43 | 0.25 |
| *Il1b* | 15 | 0.04 | 1.27 | 2.18 | 0.46 | 0.38 | 0.24 |
| *Irf9* | 15 | 0.01 | 1.84 | 2.13 | 0.47 | 0.48 | 0.29 |
| *Stat2* | 14 | 0.01 | 1.54 | 2.12 | 0.47 | 0.55 | 0.29 |
| *Gbp3* | 14 | 0.03 | 4.64 | 1.94 | 0.52 | 0.52 | 0.26 |
| *Psmb8* | 14 | 0.14 | 2.23 | 2.00 | 0.50 | 0.41 | 0.22 |
| *Psmb9* | 13 | 0.09 | 3.40 | 2.04 | 0.49 | 0.41 | 0.21 |
| *Gbp2* | 13 | 0.06 | 6.24 | 2.00 | 0.50 | 0.45 | 0.23 |
| *Dhx58* | 12 | 0.01 | 1.49 | 2.19 | 0.46 | 0.52 | 0.27 |
| *Tapbp* | 12 | 0.08 | 1.52 | 2.74 | 0.37 | 0.39 | 0.36 |

The degree distribution of nodes of the PPI network of IFN-γ signaling was analyzed and a power-law distribution was fitted to it. The power law fitting analysis exhibited the Kolmogorov-Smirnov (KS) statistic of 0.1, exponent of 4.76, coefficient of determination (R^2^) value of 1 and p-value as 0.3183. This suggests that the network possesses scale-free architecture. This indicates that most of the nodes have a small number of connections, but a few nodes have many connections. This distribution pattern also supported this notion in the plot of frequency polygon (Fig. S6A). Often, scale-free networks tend to exhibit the properties of a small-world network that is characterized by a high clustering coefficient and short average path lengths. This can be represented as L > L_random_ but C >> C_random_, where L is the characteristic path length and C is the clustering coefficient [17]. To test it, one thousand random networks were generated with the same number of nodes and edges of the IFN-γ signaling PPI network [18]. The random network analysis yielded mean C_random_ = 0.09 and mean L_random_ = 2.42. The average shortest path length of the IFN-γ signaling PPI network was 2.553, which was higher than the mean L_random_. The clustering coefficient of the PPI network of IFN-γ induced DEGs was 0.61, that was higher by 6.77 times than the mean C_random_. This result indicates that the connection topology of the PPI network of IFNγ induced DEGs exhibits small-world properties (Fig. S6B).

A total of 15 hubs were identified. These hubs, listed in descending order of degrees, were as follows: *Stat1*, *Irf7*, *Irf1*, *Cxcl10*, *Isg15*, *Il6*, *Il1b*, *Irf9*, *Stat2*, *Gbp3*, *Psmb8*, *Psmb9*, *Gbp2*, *Dhx58*, and *Tapbp*. The top 9 hubs, as determined by their degree, also exhibited the highest values in betweenness centrality (Table 2). The hubs exhibited a similar trend in correlation between degree, betweenness centrality, and log_2_ (fold change), like the overall network, where log2 fold change showed no correlation with degree and betweenness centrality, however, degree and betweenness centrality showed positive correlation (Fig. S7). The bottlenecks were *Tap1*, *Ifi204*, *Parp14* and *Ifi47*.

### Nicotinamide metabolism influences NO production and cell growth during IFN-γ-mediated responses

We investigated the expression of genes involved in NO and ROS formation in RAW 264.7 macrophages. The constitutive but low-level NO-producing genes, *Nos1* and *Nos3*, remained unaffected, with mean log_2_ (fold change) of -0.19 and -0.2, respectively. The expression of negative regulators of NO production, *Arg2* (which depletes L-arginine, the substrate for NOS2) and *Nosip* (which inhibits NOS2 by promoting its ubiquitination and degradation), was unaffected upon IFN-γ activation [19],[20]. *Cybb* gene, encoding the gp91-phox component of the phagocyte oxidase enzyme complex to produce ROS in macrophages, was markedly upregulated with a mean log_2_ (fold change) of 1.6 and p-value of ∼0.25. Importantly, *Nos2*, encoding the inducible NO-producing enzyme, responsible for producing a massive amount of NO in IFN-γ-activated macrophages, was upregulated with an average log2 (fold change) of 1.49 and a p-value of ∼0.07 (Fig. 5A).

**Figure 5.**
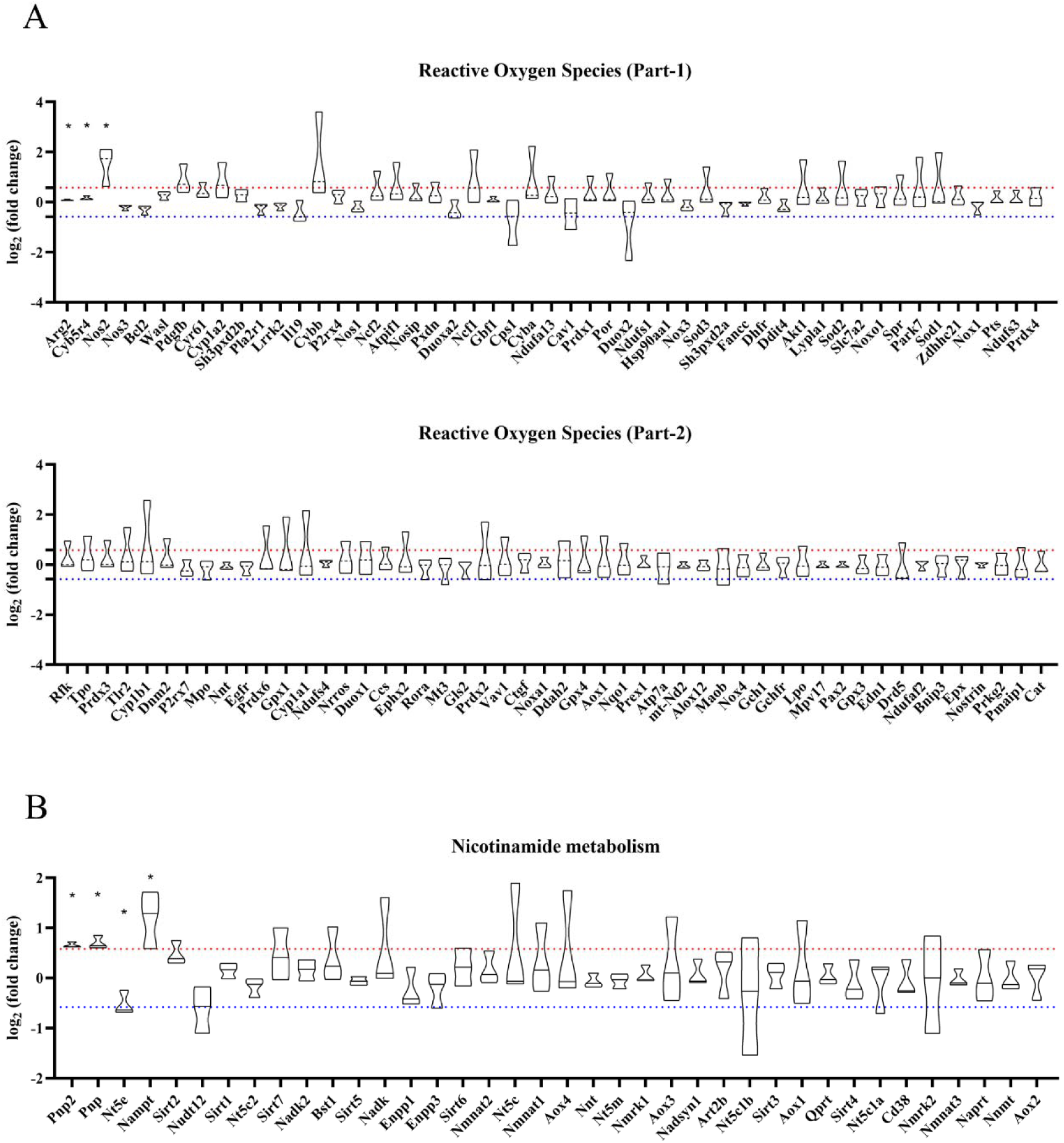
IFN-γ activation upregulates *Nos2* as a ROS generator and *Pnp*, *Pnp2* and *Nampt* as major positive regulators of nicotinamide metabolism. The expression of genes involved in ROS generation and nicotinamide metabolism is and their modulation by IFN-γ is shown.’*’ indicates a p-value < 0.05, determined by statistical comparison of log2(fold change) values using a T-statistic on Microsoft Excel (version 2404) across three independent experiments.

Among glucose transporters, *Slc2a6* (encoding Glut6) was upregulated with a mean log2 (fold change) of 0.8. *Slc2a1* (encoding Glut1) was slightly upregulated with a mean log2 (fold change) of 0.54. Among glycolytic genes, no gene was significantly upregulated. However, several genes showed a tendency towards upregulated expressions across independent experiments, notably *Hk2*, *Eno1*, and *Gapdh*. *Ldha*, involved in converting pyruvate to lactate, was upregulated with a mean log2 (fold change) of 0.66. Genes involved in the TCA cycle and electron transport system exhibited no notable expression changes (Fig. S8, S9).

Among functional enrichment of metabolic processes, metabolic reprogramming through increased NAD formation was enormously significant in the KEGG pathway and biological process gene ontology. The associated genes were *Nampt*, *Pnp*, and *Pnp2*, with a 7.3% association score in KEGG pathways and 100% association in the biological process, having the term p-value of less than 0.0001 with the name of nicotinamide riboside catabolic process (Fig. 3). *Nampt*, *Pnp*, and *Pnp2* had an average log2 (fold change) of 1.19, 0.7 and 0.65 respectively. These three genes were significantly upregulated with a p-value < 0.05 across independent experiments among all genes involved in NAD metabolism (Fig. 5B).

These observations indicate that IFN-γ-activated macrophages upregulated the NO and ROS-producing genes, *Nos2* and *Cybb*, and genes involved in NAD metabolism. Nicotinamide metabolism genes: *Nampt*, *Pnp*, and *Pnp2*, cumulatively have the potential to enhance glycolysis via increasing intracellular NAD+ pool to increase lactate production, and lead to metabolic reprogramming.

To test whether IFN-γ enhances intracellular NAD levels, RAW 264.7 macrophages were treated with varying doses of IFN-γ for different durations. A significant increase in NAD levels was observed 12 hours and 24 hours post-treatment in cells exposed to higher IFN-γ doses (Fig. 6A). To explore the role of NAD in IFN-γ-induced NO production, we employed pharmacological modulators. Forodesine hydrochloride, a PNP inhibitor, significantly reduced NO production and IFN-γ-induced lower cell growth in RAW 264.7 and H6 hepatoma cells (Figs. 6B, S10A). Conversely, β-NMN supplementation enhanced NO production at a suboptimal IFN-γ dose (5 U/mL) but did not increase NO levels at higher doses (20 U/mL), where NO production reached saturation. Importantly, β-NMN improved cell growth under IFN-γ treatment (Figs. 6C, S10B). In conclusion, these findings indicate that IFN-γ induces genes involved in NAD metabolism, influencing NO production and cell growth.

**Figure 6.**
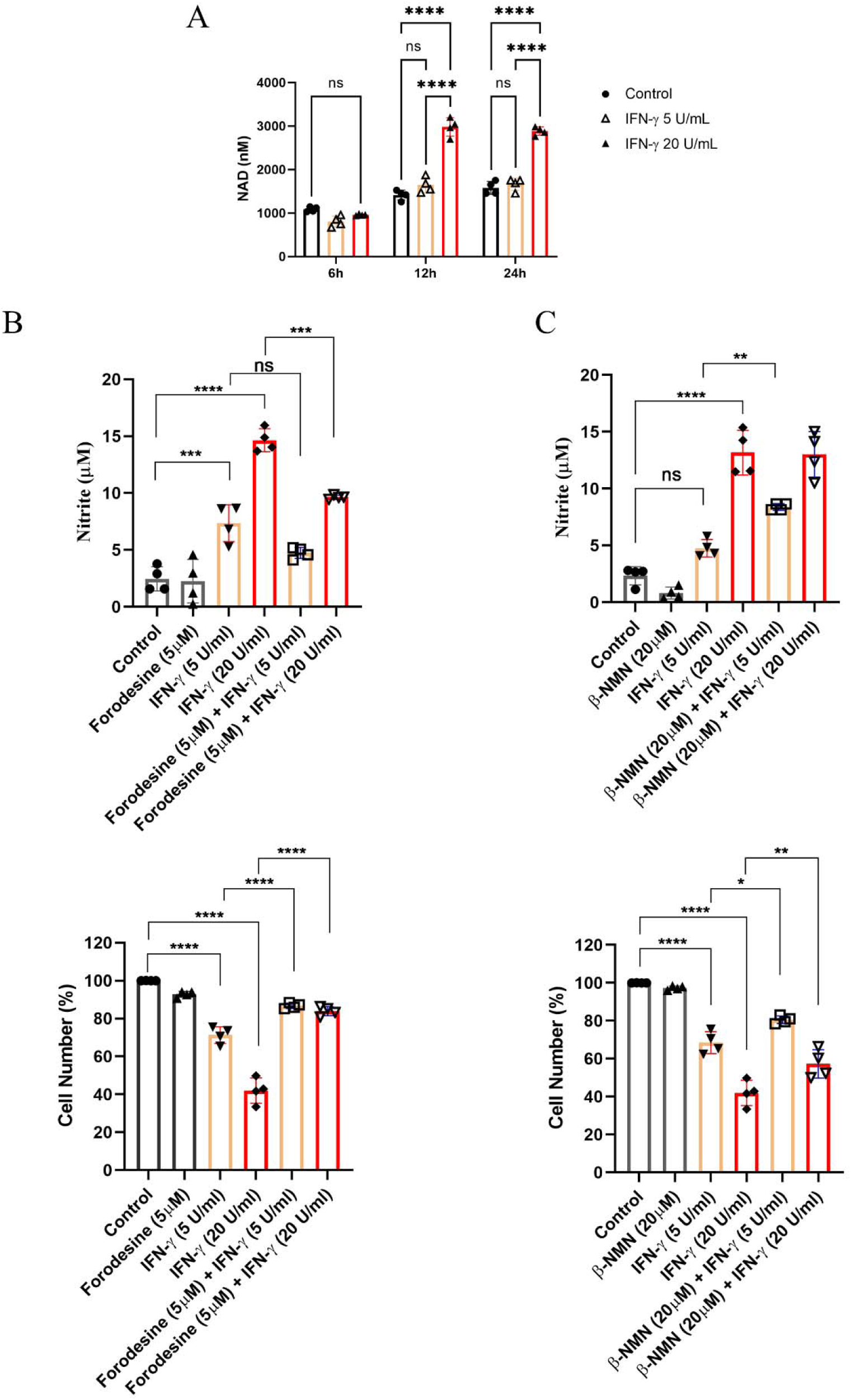
IFN-γ activation elevates intracellular NAD^+^ levels which modulate NO production in macrophages. The concentration of NAD^+^ was estimated in kinetic manner post-activation with IFN-γ (A). Forodesine (B) was used to inhibit PNP, while β-NMN supplementation (C) served as the NAD^+^ precursor to increase intracellular NAD^+^ concentration. The concentration of nitrite in cell-free supernatant was estimated using Griess assay. The percentage of cell number was measured using Trypan blue dye exclusion assay. The data were acquired after 24 hours of activation (B, C). Statistical analyses were performed using ordinary one-way ANOVA with Sidak’s multiple comparisons tests. (*), (**), (***), and (****) indicate the statistical differences of p < 0.05, p < 0.01, p < 0.001, and p < 0.0001 between the compared groups. Each data point is representative of an independent experiment. Data are represented as mean ± SD of 3-5 independent experiments.

### Gene expression analyses across healthy tissues reveal major IFN-γ-induced upregulated genes are lowly expressed in brain and highly expressed in EBV-transformed cells

Next, we focused on major IFN-γ signaling components, including the NO-dependent expression program (*Nos2*, *Klf6*, *Zfp36*), nicotinamide metabolism genes (*Nampt* and *Pnp*) etc and analyzed their expression, using data from the Genotype-Tissue Expression (GTEx) project. Under homeostatic conditions, *NOS2* exhibited the highest expression levels in the brain (specifically the cortex and frontal cortex), terminal ileum, transverse colon, pituitary gland, and lung. Many cell or tissue types demonstrated concurrent expression of *KLF6* and *ZFP36*, with expression levels above a log_10_(TPM+1) value of 2 under healthy conditions. These cell or tissue types include cultured fibroblasts, skin (both sun-exposed lower leg and non-sun-exposed suprapubic regions), arteries (coronary, tibial, and aorta), esophagus (muscularis), lung, breast (mammary tissue), adipose tissue (both subcutaneous and visceral omentum), uterus, bladder, ectocervix, fallopian tube, vagina, and tibial nerve (Fig. S11). In addition, the nicotinamide metabolism genes, *NAMPT* and *PNP*, exhibited concurrently high expression in the esophageal mucosa, visceral omentum adipose tissue, and EBV-transformed lymphocytes (Fig. S12). The lung was among the top three tissues exhibiting concurrent expression of the cytokine and chemokine hubs *CXCL10*, *IL6*, and *IL1B* under healthy conditions (Fig. S13).

EBV-transformed lymphocytes exhibited the highest expression of the IFN-γ signaling hubs: *STAT1*, *IRF7*, *PSMB8*, *PSMB9*, *ISG15*, and *DHX58*, with median log_10_(TPM+1) values exceeding 2 for all genes except *DHX58*. *IRF1*, *CXCL10*, *IRF9*, and *TAPBP* were highly expressed in EBV-infected lymphocytes, ranking among the top five tissues for these genes. Although EBV-transformed lymphocytes were not among the top five for *STAT2* expression, they still had a median log_10_(TPM+1) value greater than 2 (Fig. 7, Fig. S14, S15, S16, S17). EBV-transformed lymphocytes, lungs, and spleen exhibited the highest expression of the antigen processing and presentation hubs *PSMB8*, *PSMB9*, and *TAPBP*, ranking among the top three cells or tissues (Fig. S17).

**Figure 7.**
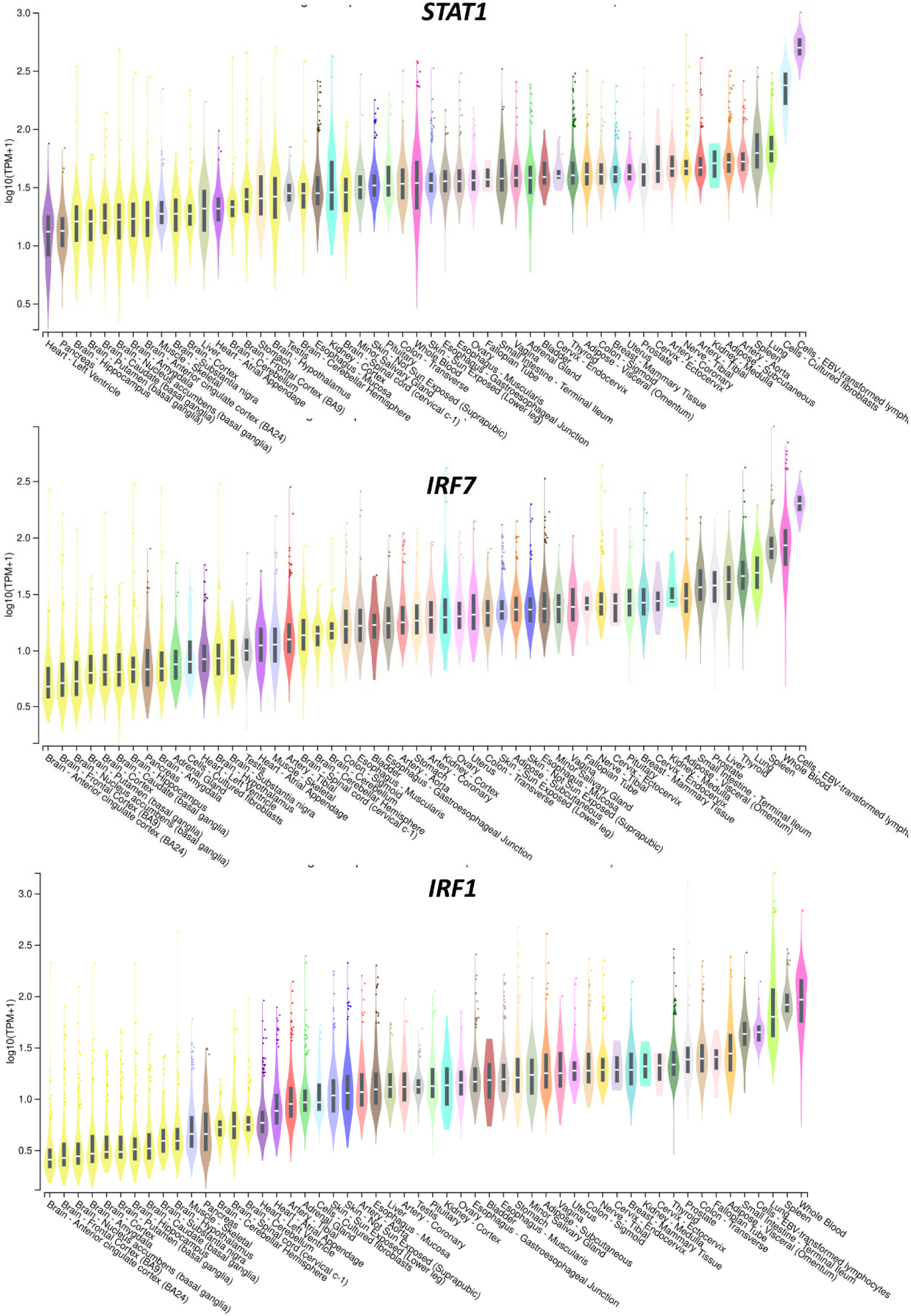
EBV-transformed lymphocytes highly express the top three hubs of the IFN-γ signaling PPI network: *STAT1*, *IRF7* and *IRF1*. The data of bulk tissue gene expression profile was obtained from GTEx Analysis Release V8. Expression values are shown in transcripts per million (TPM), calculated from a gene model with isoforms collapsed to a single gene. No other normalization steps have been applied. Box plots are shown as median and 25^th^ and 75^th^ percentiles; points are displayed as outliers if they are above or below 1.5 times the interquartile range. Color assigned to each tissue.

Furthermore, several genes demonstrated high expression in other tissues: *IRF9* in the spleen; *STAT2* in the pituitary, endocervix, nerve, and spleen; and *GBP2* in whole blood. All these tissues showed median log10(TPM+1) values greater than 2 for the respective genes (Fig. S14, S15). Notably, nicotinamide metabolism genes, NOS-dependent *Klf6* and *Zfp36*, and all the hubs demonstrated low expression across components from healthy brain, which may be important with regards to the immunological landscape in brain (Fig. 7, S11-17).

### Gene-disease association analysis reveals major IFN-γ-induced upregulated genes are involved in multiple immunopathological conditions

IFN-γ-induced significantly upregulated genes were input in the DisGeNET plug-in of Cytoscape to identify diseases associated with genes of IFN-γ signaling. The significant gene-disease associations revealed diseases ranging from immunodeficiencies to inflammatory diseases, autoimmune diseases, and cancer. The most significant hub gene of IFN-γ signaling network, *STAT1*, emerged to be associated with immunodeficiency conditions of familial candidiasis, mycobacteriosis and viral infections (Fig 8). Furthermore, *STAT3* was implicated in Job syndrome, characterized by elevated IgE levels, and infantile-onset multisystem autoimmune disease, manifesting as a spectrum of autoimmune disorders affecting multiple organs. Additionally, *IRF8* was linked to immunodeficiency conditions involving monocyte and dendritic cell deficiencies, as well as mycobacteriosis. The components of the antigen processing and presentation: *TAP1*, *TAP2* and *TAPBP* demonstrated associations with type 1 bare lymphocyte syndrome; and PSMB8 with proteasome-associated autoinflammatory syndrome. Moreover, the upregulated expression of cytokine expressing genes induced by IFN-γ, such as *IL6*, exhibited associations with diverse pathological conditions, including depressive disorders and congestive heart failure. On the other hand, *IL1B* was notably associated with mental depression, myocardial infarction, hyperalgesia, and ulcerative colitis. Further downstream in the IFN-γ signaling pathway, several genes were found to be associated with various diseases, notably, *ICAM1* with diabetic retinopathy, *RNF213* with Moyamoya disease, *PNP* with purine-nucleoside phosphorylase deficiency, *CD40* with type 3 hyper-IgM immunodeficiency syndrome and *ISG15* with immunodeficiency leading to basal ganglia calcification. Apart from the immunological disorders, some key upregulated genes of IFN-γ signaling demonstrated associations with malignant neoplasms. *STAT3* was associated with T-cell Large Granular Lymphocyte Leukemia, presenting autoimmune manifestations, while *IRF1* and *NOS2* exhibited associations with malignant neoplasms of the stomach and breast, respectively.

**Figure 8.**
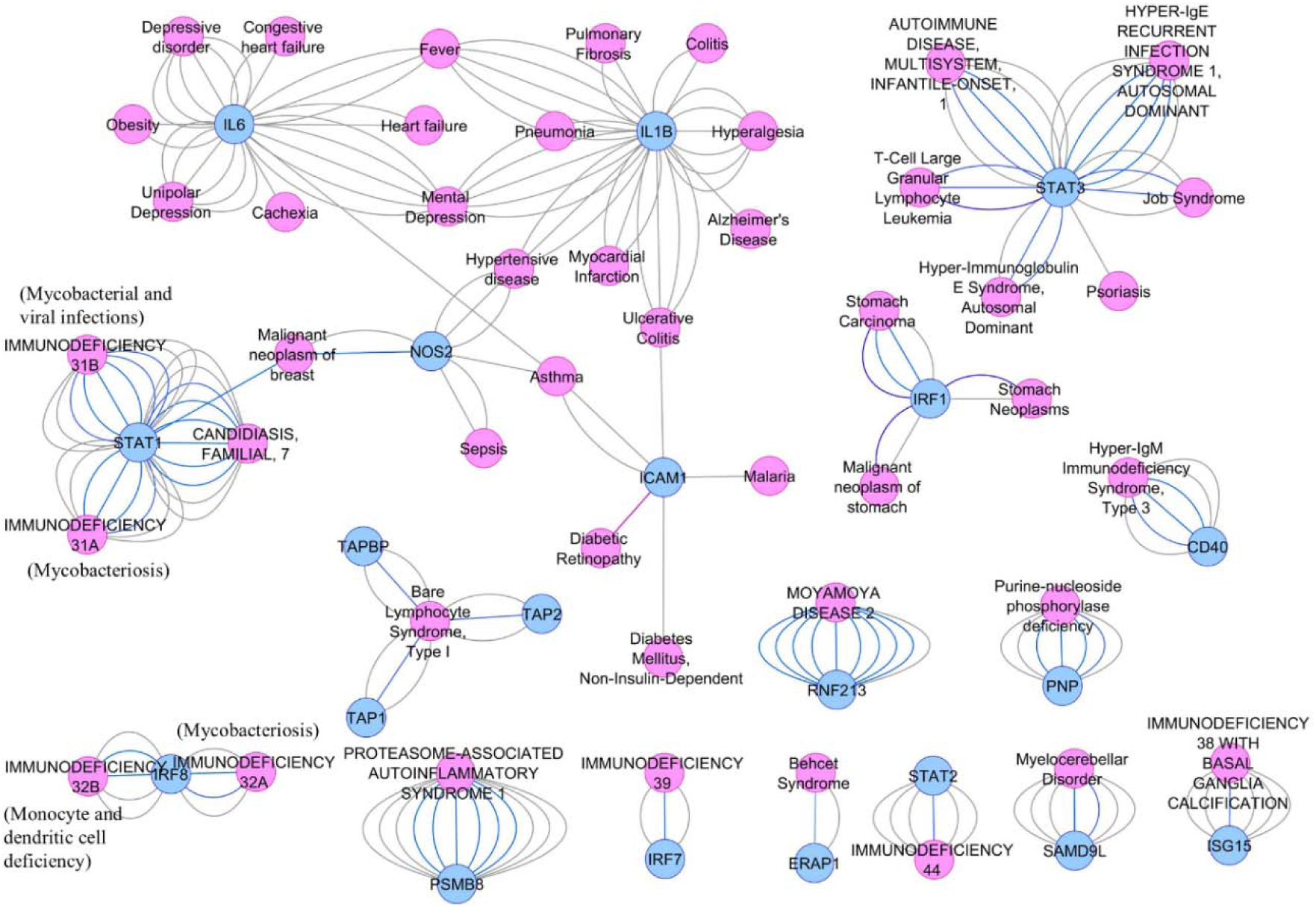
DisGeNET analysis reveals gene-disease associations of upregulated DEGs of IFN-γ signaling with human diseases. The DisGeNET network was constructed from curated sources with a minimum cut-off score of 0.6. Edge stroke colors indicate as follows: grey for biomarker, pink for causal mutation, blue for genetic variation, and purple for germline causal mutation.

Next, we focused on diseases associated with multiple genes. The diseases and their corresponding genes are as follows: asthma (*ICAM1*, *IL6*, *NOS2*), fever (*IL6*, *IL1B*), mental disorder (*IL6*, *IL1B*), hypertensive disease (*IL1B*, *NOS2*), malignant neoplasm of the breast (*STAT1*, *NOS2*), and ulcerative colitis (*ICAM1*, *IL1B*). Among the genes studied, *IL1B* was connected to the highest number of diseases implicated in four distinct conditions, followed by *IL6* and *NOS2*, each associated with three diseases. *ICAM1* was linked to two diseases, while *STAT1* was associated with one disease (Fig 8). We identified key information after reviewing the research articles demonstrating gene-disease associations on DisGeNET with these genes. *IL1B* and *IL6*, both pro-inflammatory signaling molecules, were implicated in triggering and perpetuating fever, mental disorders, hypertensive disease, and ulcerative colitis. *NOS2* was found to influence endothelial cell functions in asthma and hypertensive disease, and it played a significant role in promoting aggression in breast cancer. *STAT1*, through its involvement in IFN-γ signaling, was associated with an anti-tumor response in breast cancer. *ICAM1*, a critical cell adhesion molecule, showed elevated expression in asthma and ulcerative colitis. Notably, genetic polymorphisms such as rs5491 (K56M) and rs5498 (K469E) in *ICAM1* were linked to an increased risk of childhood asthma, particularly upon environmental tobacco exposure (Table 3). Our Comprehensive analysis underscores the multifaceted roles of these genes in various diseases.

**Table 3.** IFN-γ-inducible genes which are connected to more than one disease on DisGeNET analysis. The numbers in the table denote PMIDs with hyperlink. ‘-’ indicates not associated according to DisGeNET output.

| Names | *IL1B* | *IL6* | *NOS2* | *ICAM1* | *STAT1* |
| --- | --- | --- | --- | --- | --- |
| Asthma | - | 29902480 | 19800904 | 25003170<br>17014439 | - |
| Fever | 11852909<br>22143887<br>9952427<br>25164664<br>15384034 | 11852909<br>7537110<br>15384034 | - | - | - |
| Hypertensive disease | 27292124<br>27847271<br>27659729 | - | 25101153<br>27292124<br>18605955 | - | - |
| Malignant neoplasm of breast | - | - | 15631943<br>37169743 | - | 38354509 |
| Mental depression | 25151093<br>25315113<br>26002078<br>25286107<br>24218043 | 23813968<br>25350786<br>25903726<br>25760924<br>24491956 | - | - | - |
| Ulcerative colitis | 15955209<br>22119283<br>20452301<br>12133438 | - | - | 15553846 | - |

## Discussion

The canonical IFN-γ signaling pathway entails Janus Kinase (JAK)-Signal Transducer and Activator of Transcription (STAT) activation, leading to the transcription of primary response genes (e.g., *Irf1*, *Cxcl9*, *Cxcl10*, *Nos2*) containing *Gamma Activating Sequences*. Phosphorylated homodimerized STAT-1α mediates this process. Subsequently, transcription factors derived from the primary response genes induce the transcription of secondary response genes (e.g., *Ccl2*, *Pim1*, *Fn1*) [21]. An upregulated *Nos2* transcription is regulated through participation of different second messengers – JAK2, STAT-1α, MAP kinase kinase (MEK1/2), extracellular signal-regulated kinases 1 and 2 (Erk1/Erk2), HIF-1α, and nuclear factor kappa B (NF-κB), leading to elevated NO formation in macrophages [21]. Our study aimed to investigate the potential influence of nitric oxide (NO) on the impact of IFN-γ-induced gene expression. Through analysis of the global transcriptome following IFN-γ activation and NOS inhibition via LNMA at the early 6-hour timepoint revealed that NOS activity exerts regulatory control over a limited subset of genes with notable effects observed on *Klf6* and *Zfp36*. Furthermore, our exploration of TCGA datasets pertaining to various cancers suggested a weak correlation between the expression of these genes and Nos2, potentially due to variances in NOS2 regulation and functions between mice and human systems [22].

Our study reveals a novel finding indicating a strong correlation between the expression levels of *KLF6* and *ZFP36* across multiple cell types and tissues (Fig. 2C, S11). *KLF6* acts as a tumor suppressor gene, involved in regulating cancer development and progression by transactivating p21, reducing cyclin D1/cdk4 complexes, inhibiting c-Jun proto-oncoprotein activities, decreasing VEGF expression, and inducing apoptosis [23]. On the other hand, *KLF6-SV1*, an oncogenic splice variant of *KLF6*, is overexpressed in several human cancers, promoting tumor growth and dissemination in models of prostate and ovarian cancer [24]. KLF6 also contributes to myeloid cell plasticity in the pathogenesis of inflammatory bowel disease, where it promotes pro-inflammatory gene expression through NF-κB signaling and suppresses anti-inflammatory gene expression [26]. ZFP36 plays a crucial role in regulating inflammation by controlling the expression of pro-inflammatory cytokines and chemokines. It achieves this by binding to the 3’-untranslated regions of specific mRNAs encoding these proteins and promoting their degradation through a process called mRNA destabilization [27]. This function is particularly important in macrophages, where ZFP36 is known to regulate the expression of TNF-α, a key cytokine involved in inflammation. Accordingly, the loss of ZFP36 function in mice leads to a severe inflammatory phenotype characterized by myeloid hyperplasia, arthritis, failure of weight gain, and autoimmunity [28], [29]. In bone marrow, the expression of KLF6 and ZFP36 is regulated by social stress and miR-221/222 cluster and are involved in controlling myeloid and granulocytic differentiation of hematopoiesis in mice [30]. In this investigation, we identified that the expression of KLF6 and ZFP36 is regulated by NOS2 within the context of IFN-γ signaling, suggesting a potential influence on macrophage functions.

IFN-γ-activated cells can be classified into two groups: (1) non-NO-producing cells, for example, B16F10 melanoma, CT26 colon carcinoma, EL4 thymoma, Neuro2A neuroblastoma, TA3 mammary carcinoma, and WEHI-3 myelomonocytic leukemia; (2) NO-producing cells, for example, H6 hepatoma, Renca Renal adenocarcinoma, L929 fibrosarcoma and RAW 264.7 monocyte/macrophage cell lines [3 ,4, 5]. The NO-producing cells exhibit a metabolic shift towards enhanced glycolysis. In H6 cells, IFN-γ stimulation triggers an NO-dependent increase in glycolytic flux [5]. Lactate is a major immunosuppressive agent in the tumor microenvironment, inhibiting immune clearance [31]. Effective immune clearance necessitates intact IFN-γ signaling, as demonstrated by the poor response to immunotherapy observed in cancer patients with impaired IFN-γ signaling [32, 33, 34]. In this study, we report that RAW 264.7 macrophages produce NO and lactate upon IFN-γ activation in NO-dependent manner, suggesting a similar mechanism like H6 cells (Fig. 1A). Whether lactate production by tumors and macrophages upon IFN-γ activation significantly shapes the immunosuppressive tumor microenvironment and influences the outcomes of tumor-macrophage interactions, remain to be elucidated.

Studies on macrophages revealed that IFN-γ-activated M1-polarized macrophages undergo swift activation of aerobic glycolysis and heightened lactate production, concurrent with a decline in oxidative phosphorylation, bolstering cell viability and sustaining inflammatory responses. The elevated glycolytic flux plays a pivotal role in providing adenosine triphosphate (ATP), crucial for perpetuating IFN-γ induced JAK-STAT-1 signaling, characterized by STAT-1 phosphorylation, NOS2 activation, and subsequent NO and ROS production [6, 5]. Our analysis of NO-dependent early changes in IFN-γ modulation of gene expression affects only a small subset of genes. The RNA-seq data suggests a trend towards upregulation of glycolytic genes under IFN-γ activation, albeit not statistically significant (Fig. S8). However, in metabolic context, an important finding in our analyses is the functional enrichment of nicotinamide metabolism (Fig. 3, 5B). Three pathways maintain cellular NAD^+^: the de novo pathway from tryptophan, the Preiss-Handler pathway using nicotinic acid, and the salvage pathway utilizing nicotinamide riboside or nicotinamide. IFN-γ activation significantly upregulates NAD metabolism genes (*Nampt*, *Pnp*, *Pnp2*) involved in the salvage pathway, potentially elevating NAD^+^ levels. Elevated levels of NAD^+^ promote accelerated glycolysis by enhancing the activity of glyceraldehyde-3-phosphate dehydrogenase, which facilitates the conversion of glyceraldehyde-3-phosphate to 1,3-bisphosphoglycerate. As glycolysis progresses, NADH is oxidized back to NAD^+^ by lactate dehydrogenase, converting pyruvate to lactate, maintaining aerobic glycolysis, and facilitating elevated lactate production [35]. Hence, early upregulation of NAD^+^ biosynthesis genes in IFN-γ signaling likely enhances glycolytic flux, crucial for sustaining JAK-STAT pathway activity amidst low NO levels. As NOS2 accumulates and generates higher levels of NO and RNS/ROS, it inhibits mitochondrial function, possibly in synergy with NAD^+^, ensuring sustained elevations in glycolytic activity.

NAD^+^ metabolism plays a vital role in maintaining redox homeostasis by balancing oxidant and antioxidant levels [35]. IFN-γ signaling disrupts the redox balance by inducing nitric oxide (NO) production, leading to nitrosative and oxidative stress [4, 5]. Our findings demonstrate that inhibiting the NAD^+^ salvage pathway with Forodesine reduces nitrite levels and enhances cell survival (Fig. 6, S10). This impairment may result from a limited supply of essential substrates (L-arginine) and cofactors (NADPH, FAD, FMN), which are critical for the metabolically demanding process of NO production by NOS2 upon IFN-γ signaling [36]. Conversely, β-NMN supplementation restored NAD^+^ levels and boosted NOS2 activity, significantly enhancing NO production under suboptimal IFN-γ signaling. This dual effect—elevating NO levels to strengthen antimicrobial defenses while protecting cells from nitrosative and oxidative damage— highlights the therapeutic potential of modulating NAD^+^ metabolism to balance immune response and tissue protection. Although our study focused on nicotinamide metabolism within an *in vitro* model of IFN-γ activated macrophages, the implications extend beyond this setting. Nicotinamide metabolism is perturbed in various disease pathologies. For instance, dysregulation of the NAD^+^ salvage pathway occurs in many lower respiratory tract infections triggered by *Streptococcus* pneumoniae and SARS-CoV-2 infection. Also, Nicotinamide demonstrates efficacy in reducing disseminated candidiasis, and NAMPT blockade shows promise in ameliorating acute intestinal inflammation in inflammatory bowel disease [37 , 38 , 39 , 40]. These indicate the possibility of glycolytic perturbations across a spectrum of inflammatory scenarios.

Regulation of IFN-γ signaling involves a diverse array of post-translational modifications (PTMs), influencing cellular functions. Among these PTMs, phosphorylation of the IFN-γ receptor 2 (IFNGR2) facilitates its membrane translocation and interaction with IFNGR1, essential for the activation of the JAK-STAT pathway in macrophages [41]. Concurrently, ubiquitination targets IFNGR1 for proteasomal degradation, highlighting its crucial role in STAT1 activation and IFN-γ-dependent gene expression [42]. Furthermore, N-linked glycosylation of the IFN-γ receptor contributes to ligand binding and modulates receptor stability, impacting cellular sensitivity to IFN-γ signaling [43, 44]. NO mediates heme nitrosylation, impacting cytochrome c oxidase and soluble guanylate cyclase activities [45]. NO-mediated PTMs such as S-nitrosylation and tyrosine nitration affect cellular signaling with implications in inflammation and neurodegenerative diseases [46]. Additionally, ISGylation patterns differ between IFN-γ-activated ERα- and ERα+ breast cancer cells and influence cancer aggression [47]. ISGylation also modulates macrophage functions against antiviral defense and tumor responses to IFN-γ signaling [48]. Furthermore, nicotinamide metabolism orchestrates PTMs through enzymes like sirtuins and PARPs, critically regulating cellular metabolism and DNA damage repair mechanisms [49]. Understanding the intricacies of these PTMs is essential for unraveling the complexities of IFN-γ signaling and its broader physiological implications. Our exploration through RNA-seq analyses unveils the orchestration of PTMs via ISGylation and Nicotinamide metabolism in IFN-γ signaling, suggesting regulations at the transcript level. Importantly, ISGylation emerged as a functionally enriched process, underscored by the upregulation of *Isg15*, demonstrating strong connectivity as a hub within the IFN-γ signaling protein-protein interaction network (Fig. 3A, S5B). Concurrently, the enrichment of nicotinamide metabolism pathway genes in the IFN-γ-induced transcriptome hints at the intricate regulation of PTMs through sirtuins. Furthermore, given our observations on elevated lactate production in IFN-γ signaling, it is possible that lactylation may influence IFN-γ signaling. In fact, histone lactylations during lipopolysaccharide-stimulated macrophages drives inflammatory gene expression program towards homeostasis [50].

Analyzing protein-protein interaction (PPI) networks offers a comprehensive view of how genes within signaling pathways interact, shedding light on the molecular mechanisms driving cellular responses. Recent studies employing PPI network approaches, utilizing transcriptomic data, have uncovered key hubs influencing disease pathogenesis across multiple conditions, from infectious and inflammatory diseases to cancer. For instance, in SARS-CoV-2 infection, specific hubs like RPE5 in lung tissue and BRD4 and RIPK1 across multiple tissues have been identified. Similarly, in rheumatoid arthritis and inflammatory bowel disease, distinct hubs like IFNG, STAT3, NFKB1, TNFA, CD8A, JUN, and CTLA4 have been pinpointed, highlighting the specificity of network dysregulation in different diseases [51, 52, 53]. Moreover, detailed investigations into type I interferon signaling pathways have underscored their relevance in a spectrum of infectious and autoimmune diseases, while understanding type II interferon (IFN-γ) signaling remains incomplete [54]. Delving deeper into the PPI network of IFN-γ-induced upregulated genes provided insights into the nature of the signaling system through graph theory parameters. Interestingly, the lack of correlation between gene expression changes and degree or betweenness centrality emphasizes the importance of considering network topology in deciphering signaling regulation (Fig. 4B). The scale-free architecture of the IFN-γ signaling PPI network, characterized by a few highly connected hubs and numerous lowly connected nodes, mirrors biological systems where a small number of essential nodes control downstream events (Fig. S6A). The high clustering coefficient of the network suggests tightly interconnected clusters, facilitating efficient information flow (Fig. S6B). Moreover, the small-world properties of the network indicate efficient communication between distant nodes, crucial for rapid and coordinated responses to IFN-γ stimulation. Furthermore, identifying nodes with high degrees but low topological coefficient underscores their role as connectors or bridges between different network components rather than being embedded within local clusters (Table 2). These findings offer valuable insights into the organizational principles of the IFN-γ signaling network, with implications for understanding immune modulation and identifying therapeutic targets for immune-related diseases. For example, IFN-γ signaling in neurons is characterized by muted and prolonged activation, facilitating non-cytolytic viral clearance and influencing neuronal excitability. IFN-γ signaling affects neurogenesis and behavior and is implicated in neurodevelopmental and neurodegenerative disorders [55]. Our study reveals that the primary regulators of the IFN-γ signaling pathway, including many hubs and nicotinamide metabolism genes, are least expressed in the brain under homeostasis compared to other tissues and cell types (Fig. 7, S11-17). Additionally, we report the involvement of IFN-γ signaling genes, such as *IL6* and *IL1B*, in disease states affecting brain functions like thermoregulation and mental disorders (Fig. 8). These observations suggest that investigating IFN-γ signaling in the brain, from a healthy state to developing mental disorders and neurodegenerative diseases, could yield valuable insights for developing future therapeutics to prevent the progression of neurodegenerative diseases and mental disorders.

An exacerbation of the IFN-γ signaling pathway is often linked to inflammatory and autoimmune diseases such as asthma, ulcerative colitis, systemic lupus erythematosus, and sepsis. Targeting IFN-γ signaling and nitric oxide production has shown promise in managing ulcerative colitis and sepsis in mouse models [11, 16]. IFN-γ signaling is essential in developing malignancy, playing dual roles context-dependently. IFN-γ signaling is proven to be crucial in immunotherapy outcomes in cancer patients [56]. Our study reveals that the primary regulators of the IFN-γ signaling pathway are more highly expressed in the lungs and spleen compared to other tissues in a healthy state (Fig. 7). These tissues may require more IFN-γ signaling proteins under homeostasis due to involvement in immunological functions against environmental threats, which may contribute to the progression of asthma, ulcerative colitis, and other inflammatory diseases. Moreover, we found that EBV-transformed lymphocytes, which increase the risk of diseases like systemic lupus erythematosus and can lead to diffuse large B-cell lymphoma [57], also highly express most IFN-γ signaling pathway genes under homeostasis (Fig. 7, S12, S14, S16, S17). This indicates that IFN-γ signaling proteins in EBV-transformed lymphocytes may not only predispose individuals to SLE but also play a crucial role in the malignancy of lymphocytes. Further, we report that components of IFN-γ signaling, *STAT1*, and *NOS2*, are associated with aggression of breast cancer, whereas *IRF1* is associated with stomach cancer. Whether IFN-γ signaling promotes malignancy progression or serves as a defense mechanism to prevent uncontrolled proliferation remains to be investigated. Overall, our findings underscore the importance of understanding the metabolic and transcriptomic underpinnings of IFN-γ signaling (Fig. 9), which could influence health and present new therapeutic opportunities.

**Figure 9.**
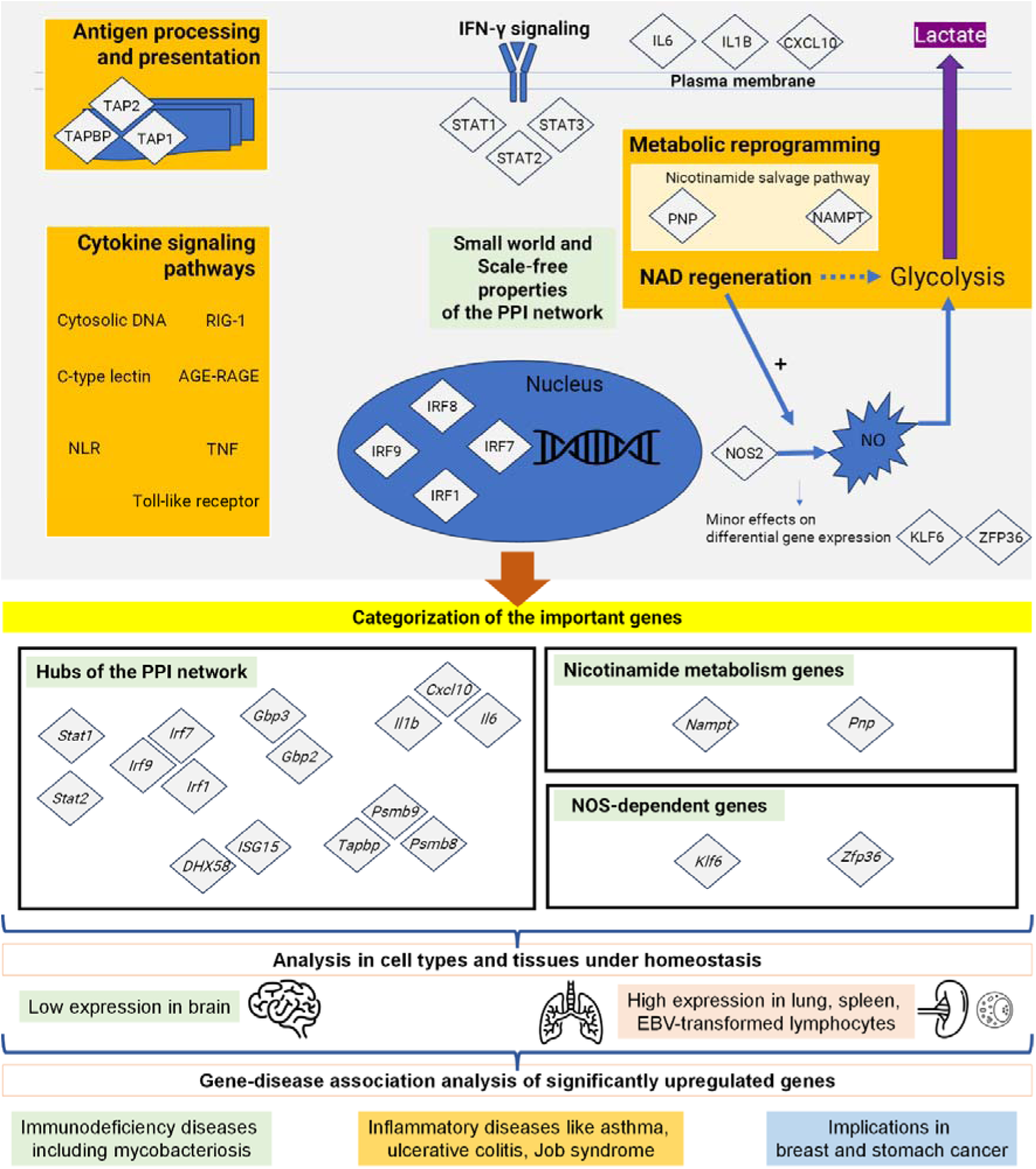
An infographic illustration demonstrates the metabolic and transcriptomic underpinnings of IFN-γ signaling in macrophages, culminating in disease pathogenesis. IFN-γ signaling in macrophages activates the canonical JAK-STAT pathway. STATs relay the signal in the nucleus to initiate transcriptional responses. IRFs get transcribed in the primary response, then initiates the secondary response transcriptional events. In our study, IFN-γ signaling transcriptome revealed the transcript level enrichment of *Stat1*, *Stat2* and *Stat3* among STATs, whereas *Irf1*, *Irf7*, *Irf8* and *Irf9* among IRFs. Among functional enrichments, antigen processing and presentation involving *Tap1*, *Tap2* and *Tapbp* is enriched. Cytokine signaling pathways including Toll-like receptor (TLR), C-type lection, TNF, NLR, RIG-1, Cytosolic DNA-sensing, AGE-RAGE are enriched. *Il6*, *Il1b* and *Cxcl10*, the genes encoding secretory cytokines, exhibit upregulated expression. Nicotinamide metabolism is highly enriched with transcript level upregulation of *Pnp* and *Nampt* genes. IFN-γ signaling leads to upregulation of *Nos2*, NO production, and NO-dependent elevation of lactate production, indicating increased glycolysis in macrophages. The enhancement of glycolysis is possibly fueled by increased NAD^+^ levels at the early stage of IFN-γ signaling with low NO levels, subsequently dominated by NO, culminating in increased lactate release. NAD^+^ contributes to optimal production of NO by NOS2. NO is potent in post-translational modifications, which might contribute to altered functions of metabolic enzymes. An analysis of NOS-dependent genes revealed that, notably, the expression of *Klf6* and *Zfp36* are upregulated by NO production. The expressions of *Klf6* and *Zfp36* are strongly correlated in many cancer types. Further, protein-protein interaction analyses revealed that IFN-γ signaling network follows scale-free and small world properties with highly dense interconnectivity. STATs (Stat1, Stat2), IRFs (Irf1, Irf7, Irf9), GBPs (Gbp2, Gbp3), cytokines (Il6, Il1b, Cxcl10), antigen processing (Psmb8, Psmb9, Tapbp), and Isg15 and Dhx58 relay the signaling responses as hubs. The major IFN-γ signaling genes from the transcriptome were categorized into hubs (important in signaling responses and promising drug targets), nicotinamide metabolism genes (relevant in metabolic reprogramming), and NO-dependent genes (relevant in cancer progression). IFN-γ signaling genes have the lowest expression in the brain and highest in lung, spleen, and EBV-transformed lymphocytes. Disease associations include immunodeficiency (mycobacteriosis), chronic inflammation (ulcerative colitis, asthma, Job syndrome), and cancers (breast, stomach).

## Materials and Methods

### Cell culture

The RAW 264.7 cell line, derived from monocytes/macrophages, was cultured in Dulbecco’s modified minimal essential medium-high glucose supplemented with 10% fetal bovine serum (FBS). The culture medium was further fortified with 5 μM β-mercaptoethanol, 100 μg/ml penicillin, 250 μg/ml streptomycin, 50 μg/ml gentamicin, and 2 mM glutamine. Cells were seeded at 3-4×10^3^ cells/mL density in 60 mm dishes and maintained at 37°C in a humidified atmosphere containing 5% CO_2_ and 95% humidity. The culture medium was refreshed every 36 hours, and cells were passaged when reaching 70-80% confluency. The cells were passaged by gentle flushing using a P1000 pipette for 2-5 minutes at room temperature.

### Nitrite, lactate and %cell number measurements

Nitrite levels, indicative of NO production, were measured using the Griess assay [58], [16]. The cell-free supernatant was mixed with distilled water in a 1:1 ratio to a total volume of 50 µL, followed by 100 µL of Griess reagent in a 96-well plate and absorbance was measured at 550 nm.

Lactate levels were measured using a colorimetric and enzymatic lactate assay kit [5]. The cell-free supernatant was diluted 1:1000 in the assay buffer. 50 µL of the reaction mixture was added to 50 µL of the diluted supernatant and incubated for 30 minutes at 37°C. The absorbance was recorded at 540 nm at 10-minute intervals using a Tecan microtiter plate reader.

Cell numbers were assessed using the Trypan Blue dye exclusion assay [3], [4], [5]. Logarithmically growing cells were seeded at an initial density of 10^4^ cells per well in a 96-well plate, treated with IFN-γ, and harvested using 0.5% Trypsin-EDTA (HiMedia, India). The cell suspension was mixed with an equal volume of 0.4% Trypan blue, and live cells were counted using a hemocytometer.

### NAD level determination

Intracellular NAD levels were estimated from IFN-γ treated cells using a colorimetric NAD/NADH estimation kit, MAK468 (Sigma Aldrich, Darmstadt, Germany). 5×10^5^ RAW 264.7 cells were seeded in each well of a 6-well plate, allowed to adhere for 12 hours, and treated with different concentrations of IFN-γ for 6, 12, and 24 hours before being subjected to the assay. Post-treatment, the supernatant was removed, and the cells were washed with cold PBS. Next, 5×10^5^ cells were collected, and processed for NAD level determination following the manufacturer’s protocol. The absorbance was recorded at 565 nm at an interval of 15 minutes using a Tecan Infinite 200 Pro microtiter plate reader (Tecan Group Ltd., Männedorf, Switzerland).

### Bulk RNA sequencing and analyses

Raw 264.7 macrophages were treated with 10 U/ml of IFN-γ for 6 hr before RNA sequencing. Bulk RNA sequencing of Raw 264.7 macrophages was performed by Clevergene (https://clevergene.in/) sequencing facility. Libraries were prepared based on polyA enrichment. The sequence data was generated using Illumina Hiseq and data quality was checked using FastQC [http://www.bioinformatics.babraham.ac.uk/projects/fastqc/] and MultiQC [59]. The data was checked for base call quality distribution, % bases above Q20, Q30, %GC, and sequencing adapter contamination (Supplementary Table 1). All the samples have passed the QC threshold (Q30>90%). The adapter sequences were ‘AGATCGGAAGAGCACACGTCTGAACTCCAGTCA’ for P7 adapter read 1 and ‘AGATCGGAAGAGCGTCGTGTAGGGAAAGAGTGT’ for P5 adapter read 2. Raw sequence reads were processed to remove adapter sequences and low-quality bases using fastp [60]. The QC passed reads were mapped onto indexed Mouse reference genome (GRCm 38.90) using STAR v2 aligner [61]. The PCR and optical duplicates were marked and removed using Picard tools [https://broadinstitute.github.io/picard/]. Gene level expression values were obtained as read counts using featureCounts software [62].

### Identification of all and NOS-dependent differentially expressed genes (DEGs)

Fold changes were calculated as the ratio of treated to control ([Treated/Control]), where treatments included IFN-γ, L-NMMA, or IFN-γ + L-NMMA. A log_2_(fold change) of 0.58 was used as the threshold for upregulated genes, while -0.58 was used for downregulated genes, provided these values met the threshold in all three independent experiments. The upregulated and downregulated genes were considered DEGs. p-values were determined from the T-statistic, which was derived from the mean log2(fold change) and the standard deviation of the mean from all three experiments. The T. DIST function was used to calculate p-values with a degree of freedom value of 2.

The identification of NO-dependent DEGs utilized Excel functions based on specific criteria across three independent experiments as follows. The ‘AND’ function isolated DEGs meeting log_2_ (fold change) thresholds of |0.58|. Conditional formatting visually distinguished DEG sets of ’Control vs. IFN-γ’ and ’Control vs. IFN-γ + LNMA’ and clubbed them together in an Excel sheet. A formula with ’IF’ and ’COUNTIF’ functions detected duplicate entries, categorizing them as NOS-independent DEGs. Further analysis identified NOS-dependent DEGs across conditions. For instance, genes upregulated in control vs. IFN-γ but unaffected or downregulated in control vs. IFN-γ+LNMA were identified. Similarly, genes showing contrasting expression profiles between the two conditions were identified. The ’OR’ function facilitated this classification by evaluating gene expression across experiments. These genes were categorized as NOS-dependent DEGs. Excel functions efficiently sorted and analyzed DEGs, revealing distinct expression patterns.

### Construction, visualization, and analysis of PPI network

PPI network: The list of significantly upregulated differentially expressed genes (DEGs) was imported in the String tool [63] of EMBL-EBI to construct the network. The minimum required interaction score was set to the medium confidence of 0.4 for the PPI network of NOS-dependent genes and highest confidence of 0.9 for IFN-γ signaling PPI network. The data was exported as TSV file and opened in Cytoscape [64]. Visualization and analysis of the network were done in Cytoscape. The hubs and bottlenecks were identified by analyzing the values of degree and betweenness centrality of the nodes in Microsoft excel. Hubs were defined as all nodes that are in the top 20% of the degree distribution. Bottlenecks were defined as the non-hub nodes that are in the top 20% in terms of betweenness centrality and connected to at least two hubs [65].

Functional enrichment analyses of KEGG pathways and Gene Ontology profile analyses were performed using ClueGO plugin of Cytoscape using the IFN-γ signaling PPI network. The marker lists were loaded as *Mus musculus* [10090]. The visual style was set at ‘significance.’ The pathways/ontologies were used as per the updated version of 25^th^ May 2022. Pathways with p-value ≤ 0.05 were displayed. The program was run on 30^th^ March 2024.

Gene-disease association network was constructed using the IFN-γ signaling PPI network in Cytoscape plug-in of DisGeNET [66]. The initial score value cut-off was set at 0.6. For gene-specific gene-disease association analyses, no such cut-off was set. The program was run on 5^th^ March 2024.

### Statistical analysis

The ordinary One-way ANOVA statistical analyses with Sidak’s multiple comparison tests were performed on GraphPad Prism software (version 8.0.2). Unless stated otherwise, the data are represented as mean + standard deviation of the mean (SD). The data distribution was tested using QQ-plot to examine the skewness. Statistical heatmap, Venn diagrams, and network biology analyses were performed on Python. Graphical representations were prepared using GraphPad Prism and Python.

## Abbreviations

DMEM: Dulbecco’s modified minimal essential medium
IFN-γ: Interferon-gamma
LNMA: NG-Methyl-L-arginine acetate salt
NO: Nitric Oxide
NOS: Nitric Oxide Synthase

## Author Contributions

AC designed experiments, performed, analyzed, and interpreted data along with writing the paper. SS performed data analyses on Python, interpreted the data, and revised the manuscript. SD performed some of the experiments and wrote parts of the manuscript. DN designed experiments, analyzed the data, revised, and approved the final manuscript.

## Acknowledgments

We acknowledge Clevergene^®^ for performing the bulk RNA-sequencing and providing us with the raw data. We greatly appreciate the support from the members of the Divisional flow cytometry facility and Central Animal Facility, IISc. The infrastructural support from DST-FIST to the Department of Biochemistry, IISc is greatly appreciated. In addition, we are thankful for the support from all members of the DpN laboratory.

## Data availability statement

The raw FASTQ files are available in GEO dataset GSE286915. Normalized expression counts, DEGs, Functional enrichment and PPI network tables are available as source data files. The GitHub link where the codes are available are as follows: https://github.com/saishyam1/IFNg_Transcriptome_Analysis_Avik_Sai.git

## References

1 Ivashkiv LB (2018) IFNγ: signalling, epigenetics and roles in immunity, metabolism, disease and cancer immunotherapy. Nat Rev Immunol 18, 545–558.

2 Förstermann U & Sessa WC (2012) Nitric oxide synthases: Regulation and function. Eur Heart J 33.

3 Prasanna SJ, Saha B & Nandi D (2007) Involvement of oxidative and nitrosative stress in modulation of gene expression and functional responses by IFNγ. Int Immunol 19, 867–879.

4 Rakshit S, Chandrasekar BS, Saha B, Victor ES, Majumdar S & Nandi D (2014) Interferon-gamma induced cell death: Regulation and contributions of nitric oxide, cJun N-terminal kinase, reactive oxygen species and peroxynitrite. Biochim Biophys Acta Mol Cell Res 1843, 2645–2661.

5 Chattopadhyay A, Jagdish S, Karhale AK, Ramteke NS, Zaib A & Nandi D (2023) IFN-γ lowers tumor growth by increasing glycolysis and lactate production in a nitric oxide-dependent manner: implications for cancer immunotherapy. Front Immunol 14, 1282653.

6 Wang F, Zhang S, Jeon R, Vuckovic I, Jiang X, Lerman A, Folmes CD, Dzeja PD & Herrmann J (2018) Interferon Gamma Induces Reversible Metabolic Reprogramming of M1 Macrophages to Sustain Cell Viability and Pro-Inflammatory Activity. EBioMedicine 30, 303–316.

7 Palmieri EM, McGinity C, Wink DA & McVicar DW (2020) Nitric oxide in macrophage immunometabolism: Hiding in plain sight. Metabolites 10, 1–34.

8 Chandrasekar BS, Yadav S, Victor ES, Majumdar S, Deobagkar-Lele M, Wadhwa N, Podder S, Das M & Nandi D (2015) Interferon-gamma and nitric oxide synthase 2 mediate the aggregation of resident adherent peritoneal exudate cells: Implications for the host response to pathogens. PLoS One 10, e0128301.

9 Henard CA & Vázquez-Torres A (2011) Nitric oxide and salmonella pathogenesis. Front Microbiol 2, 84

10 Herbst S, Schaible UE & Schneider BE (2011) Interferon gamma activated macrophages kill mycobacteria by nitric oxide induced apoptosis. PLoS One 6, e19105.

11 Yadav S, Pathak S, Sarikhani M, Majumdar S, Ray S, Chandrasekar BS, Adiga V, Sundaresan NR & Nandi D (2018) Nitric oxide synthase 2 enhances the survival of mice during Salmonella Typhimurium infection-induced sepsis by increasing reactive oxygen species, inflammatory cytokines and recruitment of neutrophils to the peritoneal cavity. Free Radic Biol Med 116, 73–87.

12 Chattopadhyay A, Joseph JP, Shyam S & Nandi D (2022) Characterizing Salmonella Typhimurium-induced Septic Peritonitis in Mice. Journal of Visualized Experiments 185, e63695.

13 Tandon M, Wu W, Moore K, Winchester S, Tu Y-P, Miller C, Kodgule R, Pendse A, Rangwala S, Joshi S & Group S (2022) SARS-CoV-2 accelerated clearance using a novel nitric oxide nasal spray (NONS) treatment: A randomized trial, The Lancet Regional Health-Southeast Asia, 3.

14 Xiao S, Yuan Z & Huang Y (2023) The Potential Role of Nitric Oxide as a Therapeutic Agent against SARS-CoV-2 Infection. Int J Mol Sci 24, 17162.

15 Zhang H, Zhang C, Hua W & Chen J (2024) Saying no to SARS-CoV-2: the potential of nitric oxide in the treatment of COVID-19 pneumonia. Med Gas Res 14, 39–47.

16 Chattopadhyay A, Joseph JP, Jagdish S, Chaudhuri S, Ramteke NS, Karhale AK, Waturuocha U, Saini DK & Nandi D (2023) High throughput screening identifies auranofin and pentamidine as potent compounds that lower IFN-γ-induced Nitric Oxide and inflammatory responses in mice: DSS-induced colitis and Salmonella Typhimurium-induced sepsis. Int Immunopharmacol 122, 110569.

17 Watts, D., Strogatz, S. Collective dynamics of ‘small-world’ networks. Nature 393, 440–442.

18 Killick KE, Magee DA, Park SDE, Taraktsoglou M, Browne JA, Conlon KM, Nalpas NC, Gormley E, Gordon S V., MacHugh DE & Hokamp K (2014) Key hub and bottleneck genes differentiate the macrophage response to virulent and attenuated Mycobacterium bovis. Front Immunol 5, 422.

19 Durante W, Johnson FK, Johnson RA. Arginase: a critical regulator of nitric oxide synthesis and vascular function. Clin Exp Pharmacol Physiol. 34, 906–11.

20 Mukherjee R, Singh DK, Patra R, et al. A novel polymorphism in nitric oxide synthase interacting protein (NOSIP) modulates nitric oxide and mortality in Human Sepsis. bioRxiv.

21 Saha B, Jyothi Prasanna S, Chandrasekar B & Nandi D (2010) Gene modulation and immunoregulatory roles of Interferonγ. Cytokine 50, 1–14.

22 Blanchette J, Jaramillo M & Olivier M Signalling events involved in interferon-c-inducible macrophage nitric oxide generation. Immunology. 108, 513–22.

23 Hoos MD, Vitek MP, Ridnour LA, Wilson J, Jansen M, Everhart A, Wink DA & Colton CA (2014) The impact of human and mouse differences in NOS2 gene expression on the brain’s redox and immune environment. Mol Neurodegener 9, 50.

24 Sabatino ME, Castellaro A, Racca AC, Carbajosa González S, Pansa MF, Soria G & Bocco JL (2019) Krüppel-Like Factor 6 Is Required for Oxidative and Oncogene-Induced Cellular Senescence. Front Cell Dev Biol 7.

25 DiFeo A, Martignetti JA & Narla G (2009) The role of KLF6 and its splice variants in cancer therapy. Drug Resistance Updates 12, 1–7.

26 Goodman WA, Omenetti S, Date D, Di Martino L, De Salvo C, Kim GD, Chowdhry S, Bamias G, Cominelli F, Pizarro TT & Mahabeleshwar GH (2016) KLF6 contributes to myeloid cell plasticity in the pathogenesis of intestinal inflammation. Mucosal Immunol 9, 1250–1262.

27 Moore M, Blachere N, Fak J, Park C, Sawicka K, Parveen S, Zucker-Scharff I, Moltedo B, Rudensky A & Darnell R (2018) ZFP36 RNA-binding proteins restrain T cell activation and anti-viral immunity. 7, e33057.

28 Zhang H, Taylor WR, Joseph G, Caracciolo V, Gonzales DM, Sidell N, Seli E, Blackshear PJ & Kallen CB (2013) MRNA-binding protein ZFP36 is expressed in atherosclerotic lesions and reduces inflammation in aortic endothelial cells. Arterioscler Thromb Vasc Biol 33, 1212–1220.

29 Snyder BL, Huang R, Burkholder AB, Donahue DR, Mahler BW, Bortner CD, Lai WS & Blackshear PJ (2024) Synergistic roles of tristetraprolin family members in myeloid cells in the control of inflammation. Life Sci Alliance 7.

30 Jani PK, Petkau G, Kawano Y, Klemm U, Guerra GM, Heinz GA, Heinrich F, Durek P, Mashreghi MF & Melchers F (2023) The miR-221/222 cluster regulates hematopoietic stem cell quiescence and multipotency by suppressing both Fos/AP-1/ IEG pathway activation and stress-like differentiation to granulocytes. PLoS Biol 21, e3002015.

31 Hayes C, Donohoe CL, Davern M & Donlon NE (2021) The oncogenic and clinical implications of lactate induced immunosuppression in the tumour microenvironment. Cancer Lett 500, 75–86.

32 Minn AJ & Wherry EJ (2016) Combination Cancer Therapies with Immune Checkpoint Blockade: Convergence on Interferon Signaling. Cell 165, 272–275.

33 Grasso CS, Tsoi J, Onyshchenko M, Abril-Rodriguez G, Ross-Macdonald P, Wind-Rotolo M, Champhekar A, Medina E, Torrejon DY, Shin DS, Tran P, Kim YJ, Puig-Saus C, Campbell K, Vega-Crespo A, Quist M, Martignier C, Luke JJ, Wolchok JD, Johnson DB, Chmielowski B, Hodi FS, Bhatia S, Sharfman W, Urba WJ, Slingluff CL, Diab A, Haanen JBAG, Algarra SM, Pardoll DM, Anagnostou V, Topalian SL, Velculescu VE, Speiser DE, Kalbasi A & Ribas A (2020) Conserved Interferon-γ Signaling Drives Clinical Response to Immune Checkpoint Blockade Therapy in Melanoma. Cancer Cell 38, 500–515.e3.

34 Larson RC, Kann MC, Bailey SR, Haradhvala NJ, Llopis PM, Bouffard AA, Scarfó I, Leick MB, Grauwet K, Berger TR, Stewart K, Anekal PV, Jan M, Joung J, Schmidts A, Ouspenskaia T, Law T, Regev A, Getz G & Maus M V. (2022) CAR T cell killing requires the IFNγR pathway in solid but not liquid tumours. Nature 604, 563–570.

35 Xie N, Zhang L, Gao W, Huang C, Huber PE, Zhou X, Li C, Shen G & Zou B (2020) NAD+ metabolism: pathophysiologic mechanisms and therapeutic potential. Signal Transduct Target Ther 5, 227.

36 Bogdan C (2015) Nitric oxide synthase in innate and adaptive immunity: An update. Trends Immunol 36, 161–178.

37 Klabunde B, Wesener A, Bertrams W, Beinborn I, Paczia N, Surmann K, Blankenburg S, Wilhelm J, Serrania J, Knoops K, Elsayed EM, Laakmann K, Jung AL, Kirschbaum A, Hammerschmidt S, Alshaar B, Gisch N, Abu Mraheil M, Becker A, Völker U, Vollmeister E, Benedikter BJ & Schmeck B (2023) NAD+ metabolism is a key modulator of bacterial respiratory epithelial infections. Nat Commun 14, 5818.

38 Xing XR, Liao Z Bin, Tan F, Zhu ZY, Jiang Y & Cao YY (2019) Effect of nicotinamide against candida albicans. Front Microbiol 10, 595.

39 Ramana C V. (2024) Regulation of a Metabolic Gene Signature in Response to Respiratory Viruses and Type I Interferon Signaling. Journal of Molecular Pathology 5, 133–152.

40 Gerner RR, Klepsch V, Macheiner S, Arnhard K, Adolph TE, Grander C, Wieser V, Pfister A, Moser P, Hermann-Kleiter N, Baier G, Oberacher H, Tilg H & Moschen AR (2018) NAD metabolism fuels human and mouse intestinal inflammation. Gut 67, 1813–1823.

41 Xu X, Xu J, Wu J, Hu Y, Han Y, Gu Y, Zhao K, Zhang Q, Liu X, Liu J, Liu B & Cao X (2018) Phosphorylation-Mediated IFN-γR2 Membrane Translocation Is Required to Activate Macrophage Innate Response. Cell 175, 1336–1351.e17.

42 Londino JD, Gulick DL, Lear TB, Suber TL, Weathington NM, Masa LS, Chen BB & Mallampalli RK (2017) Post-translational modification of the interferon-gamma receptor alters its stability and signaling. Biochemical Journal 474, 3543–3557.

43 Fischer T, Thoma B, Scheurich P & Pfizenmaier K (1990) Glycosylation of the human interferon-γ receptor: N-linked carbohydrates contribute to structural heterogeneity and are required for ligand binding. Journal of Biological Chemistry 265, 1710–1717.

44 Mo X, Kazmi HR, Preston-Alp S, Zhou B & Zaidi MR (2022) Interferon-gamma induces melanogenesis via post-translational regulation of tyrosinase. Pigment Cell Melanoma Res 35, 342–355.

45 Martí MA, Capece L, Crespo A, Doctorovich F & Estrin DA (2005) Nitric oxide interaction with cytochrome c′ and its relevance to guanylate cyclase. Why does the iron histidine bond break? J Am Chem Soc 127, 7721–7728.

46 Leon L, Jeannin JF & Bettaieb A (2008) Post-translational modifications induced by nitric oxide (NO): Implication in cancer cells apoptosis. Nitric Oxide 19, 77–83.

47 Tecalco-Cruz AC & Cruz-Ramos E (2018) Protein ISGylation and free ISG15 levels are increased by interferon gamma in breast cancer cells. Biochem Biophys Res Commun 499, 973–978.

48 Munnur D, Teo Q, Eggermont D, Lee HHY, Thery F, Ho J, van Leur SW, Ng WWS, Siu LYL, Beling A, Ploegh H, Pinto-Fernandez A, Damianou A, Kessler B, Impens F, Mok CKP & Sanyal S (2021) Altered ISGylation drives aberrant macrophage-dependent immune responses during SARS-CoV-2 infection. Nat Immunol 22, 1416–1427.

49 Munk SHN, Merchut-Maya JM, Adelantado Rubio A, Hall A, Pappas G, Milletti G, Lee MH, Johnsen LG, Guldberg P, Bartek J & Maya-Mendoza A (2023) NAD+ regulates nucleotide metabolism and genomic DNA replication. Nat Cell Biol 25, 1774–1786.

50 Zhang D, Tang Z, Huang H, Zhou G, Cui C, Weng Y, Liu W, Kim S, Lee S, Perez-Neut M, Ding J, Czyz D, Hu R, Ye Z, He M, Zheng YG, Shuman HA, Dai L, Ren B, Roeder RG, Becker L & Zhao Y (2019) Metabolic regulation of gene expression by histone lactylation. Nature 574, 575–580.

51 Asadzadeh-Aghdaee H, Shahrokh S, Norouzinia M, Hosseini M, Keramatinia A, Jamalan M, Naghibzadeh B, Sadeghi A, Sherafat SJ & Zali MR (2016) Introduction of inflammatory bowel disease biomarkers panel using protein-protein interaction (PPI) network analysis. Gastroenterol Hepatol Bed Bench 9, S8.

52 Eskandarzade N, Ghorbani A, Samarfard S, Diaz J, Guzzi PH, Fariborzi N, Tahmasebi A & Izadpanah K (2022) Network for network concept offers new insights into host-SARS-CoV-2 protein interactions and potential novel targets for developing antiviral drugs. Comput Biol Med 146, 105575.

53 Yuan S-G, Zheng K, Chen M & Zhang Z (2024) Analysis of Enrichment Pathway, Hub Gene, and Protein-Protein Interaction Network in Rheumatoid Arthritis and Construction of Molecular Subtypes in Peripheral Blood. Altern Ther Health Med 30.

54 Kerr CH, Skinnider MA, Andrews DDT, Madero AM, Chan QWT, Stacey RG, Stoynov N, Jan E & Foster LJ (2020) Dynamic rewiring of the human interactome by interferon signaling. Genome Biol 21, 1–36

55 Clark DN, Begg LR & Filiano AJ (2022) Unique aspects of IFN-γ/STAT1 signaling in neurons. Immunol Rev 311, 187–204.

56 Zaidi MR (2019) The Interferon-Gamma Paradox in Cancer. Journal of Interferon and Cytokine Research 39, 30–38.

57 Zhu QY (2023) Bioinformatics analysis of the pathogenic link between Epstein-Barr virus infection, systemic lupus erythematosus and diffuse large B cell lymphoma. Sci Rep 13, 6310.

58 Malu S, Srinivasan S, Kumar Maiti P, Rajagopal D, John B & Nandi D IFN-g bioassay: development of a sensitive method by measuring nitric oxide production by peritoneal exudate cells from C57BL/6 mice. J Immunol Methods 272, 55–65.

59 Ewels P, Magnusson M, Lundin S & Käller M (2016) MultiQC: Summarize analysis results for multiple tools and samples in a single report. Bioinformatics 32, 3047–3048.

60 Chen S, Zhou Y, Chen Y & Gu J (2018) Fastp: An ultra-fast all-in-one FASTQ preprocessor. In Bioinformatics 34, i884–i890.

61 Dobin A, Davis CA, Schlesinger F, Drenkow J, Zaleski C, Jha S, Batut P, Chaisson M & Gingeras TR (2013) STAR: Ultrafast universal RNA-seq aligner. Bioinformatics 29, 15–21.

62 Liao Y, Smyth GK & Shi W (2014) FeatureCounts: An efficient general purpose program for assigning sequence reads to genomic features. Bioinformatics 30, 923–930.

63 Szklarczyk D, Gable AL, Lyon D, Junge A, Wyder S, Huerta-Cepas J, Simonovic M, Doncheva NT, Morris JH, Bork P, Jensen LJ & Von Mering C (2019) STRING v11: Protein-protein association networks with increased coverage, supporting functional discovery in genome-wide experimental datasets. Nucleic Acids Res 47, D607–D613.

64 Shannon P, Markiel A, Ozier O, Baliga NS, Wang JT, Ramage D, Amin N, Schwikowski B & Ideker T (2003) Cytoscape: A software Environment for integrated models of biomolecular interaction networks. Genome Res 13, 2498–2504.

65 Yu H, Kim PM, Sprecher E, Trifonov V & Gerstein M (2007) The importance of bottlenecks in protein networks: Correlation with gene essentiality and expression dynamics. PLoS Comput Biol 3, 713–720.

66 Piñero J, Saüch J, Sanz F & Furlong LI (2021) The DisGeNET cytoscape app: Exploring and visualizing disease genomics data. Comput Struct Biotechnol J 19, 2960–2967.

