## Supplemental Information for "Global transcriptome analysis identifies nicotinamide metabolism to play key roles in IFN-γ and nitric oxide modulated responses"

**Supplementary materials**

**Supplementary Tables**

**Supplementary Table 1. Summary of raw sequence data and quality**

| Sample | Number of reads | Read length | GC% | %Bases > Q20 | %Bases > Q30 |
| --- | --- | --- | --- | --- | --- |
| Control-1 | 61742754 | 150 | 50 | 99.69 | 93.11 |
| Control-2 | 25847290 | 150 | 46 | 99.16 | 91.77 |
| Control-3 | 34056124 | 150 | 44 | 99.4 | 91.51 |
| IFN-γ-1 | 69672146 | 150 | 51 | 99.71 | 92.50 |
| IFN-γ-2 | 27531694 | 150 | 46.5 | 99.27 | 90.09 |
| IFN-γ-3 | 27203676 | 150 | 47 | 99.29 | 92.35 |
| LNMA-1 | 48890180 | 150 | 45 | 99.74 | 88.70 |
| LNMA-2 | 29348716 | 150 | 46 | 99.05 | 87.69 |
| LNMA-3 | 30049078 | 150 | 47.5 | 99.13 | 91.44 |
| IFN-γ+LNMA-1 | 57488492 | 150 | 50 | 99.67 | 93.64 |
| IFN-γ+LNMA-2 | 40429648 | 150 | 49 | 99.57 | 57.45 |
| IFN-γ+LNMA-3 | 29163142 | 150 | 46.5 | 99.03 | 88.77 |

**Supplementary Table 2. Read alignment statistics with number of expressed genes**

Out of a total of 52,636 genes, the numbers of expressed genes are listed.

| Sample | Reads  after QC | Mapped  reads | Mapped  reads% | Uniquely  mapped  reads | Uniquely  mapped  reads % | Unmapped  Reads | Unmapped  Reads% | No. of expressed genes |
| --- | --- | --- | --- | --- | --- | --- | --- | --- |
| Control-1 | 58294302 | 58069448 | 99.61 | 38966998 | 66.85 | 224854 | 0.39 | 34971 |
| Control-2 | 23537956 | 21138086 | 89.8 | 13697056 | 58.19 | 2399870 | 10.2 | 45721 |
| Control-3 | 31216932 | 28595404 | 91.6 | 19034060 | 60.97 | 2621528 | 8.4 | 47249 |
| IFN-γ-1 | 65426994 | 65156540 | 99.59 | 43502480 | 66.49 | 270454 | 0.41 | 35370 |
| IFN-γ-2 | 24807538 | 22261816 | 89.74 | 14396662 | 58.03 | 2545722 | 10.26 | 46075 |
| IFN-γ-3 | 25089668 | 22895102 | 91.25 | 14573682 | 58.09 | 2194566 | 8.75 | 45583 |
| LNMA-1 | 43282108 | 42807670 | 98.9 | 26655624 | 61.59 | 474438 | 1.1 | 37946 |
| LNMA-2 | 25673586 | 22982912 | 89.52 | 15290292 | 59.56 | 2690674 | 10.48 | 46488 |
| LNMA-3 | 27537408 | 23919426 | 86.86 | 15689020 | 56.97 | 3617982 | 13.14 | 47328 |
| IFN-γ+LNMA-1 | 54631846 | 54434932 | 99.64 | 36766402 | 67.3 | 196914 | 0.36 | 34428 |
| IFN-γ+LNMA-2 | 24627308 | 21958254 | 89.16 | 14218540 | 57.73 | 2669054 | 10.84 | 46467 |
| IFN-γ+LNMA-3 | 25683846 | 22465864 | 87.47 | 15106820 | 58.82 | 3217982 | 12.53 | 46497 |

**Supplementary figures:**

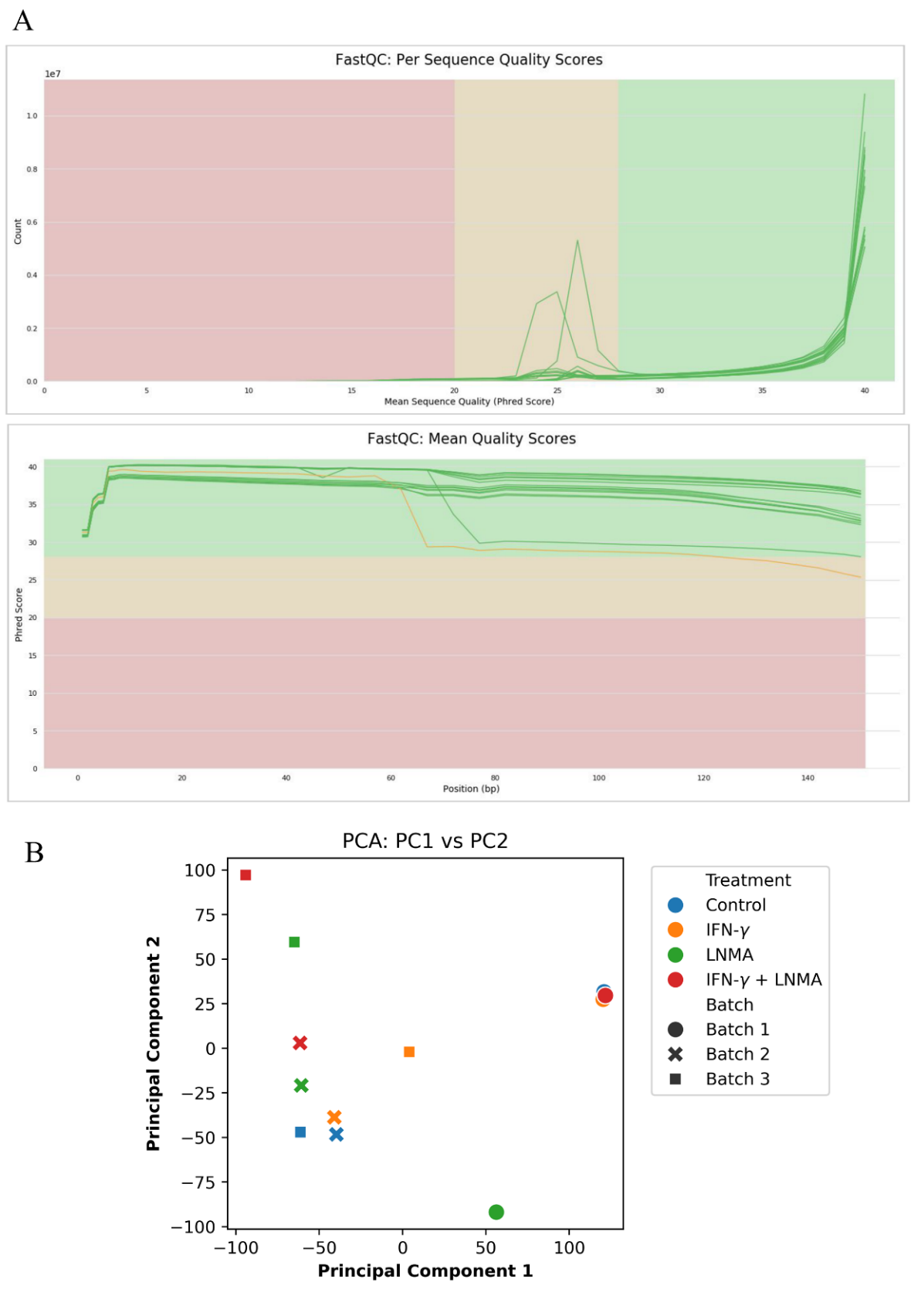

**Fig.S1. Quality assessment of reveals a good quality the RNA-seq data for downstream analyses**

Per sequence quality scores and mean quality scores of the FastQC (A). Principal Component Analysis showing the similarities and differences between samples (B).

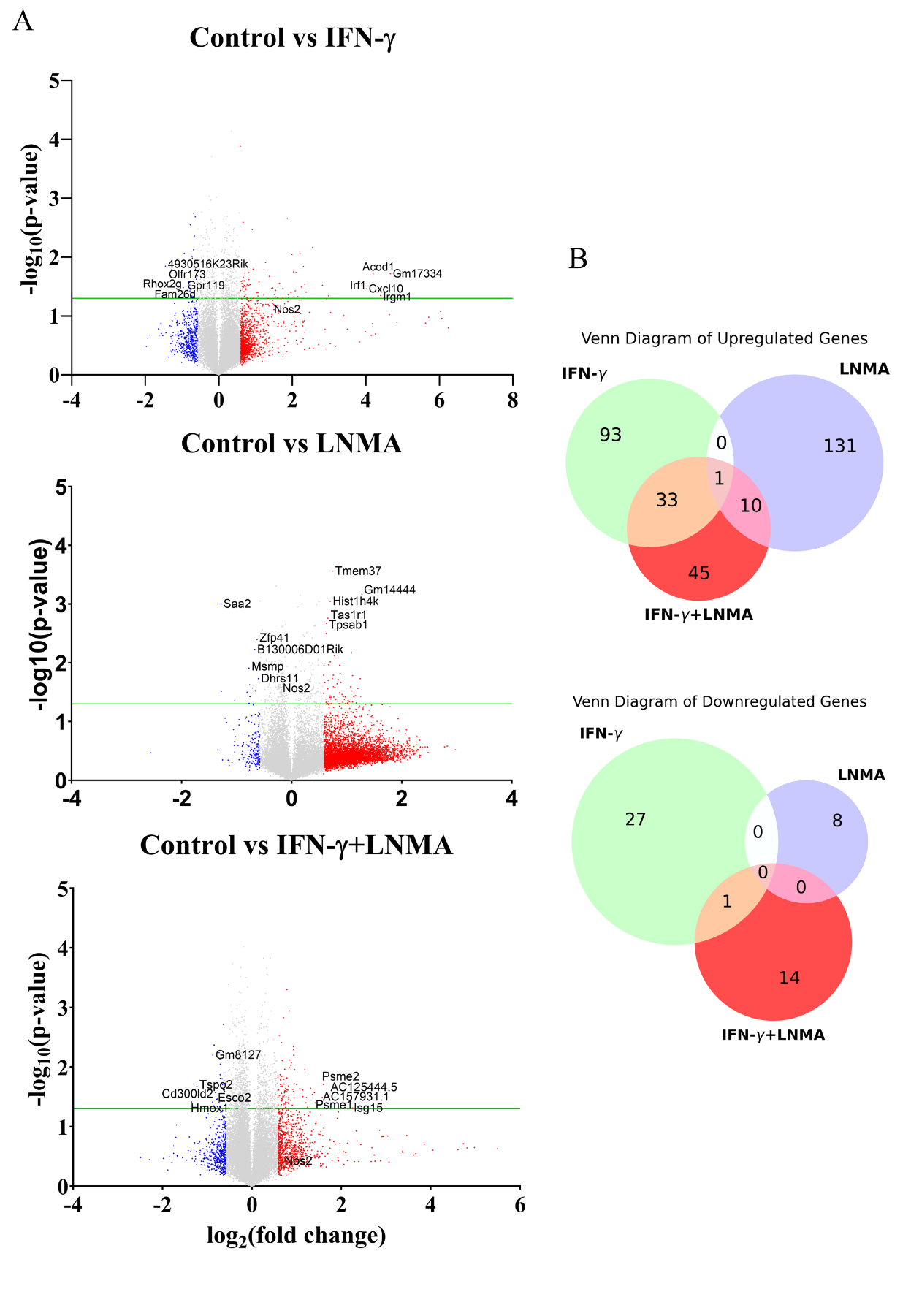

**Fig. S2. Distribution of DEGs across comparisons**

Volcano plot representation displaying significantly upregulated and downregulated genes across comparisons (A). Venn diagram displaying the number of genes shared across comparisons (B).

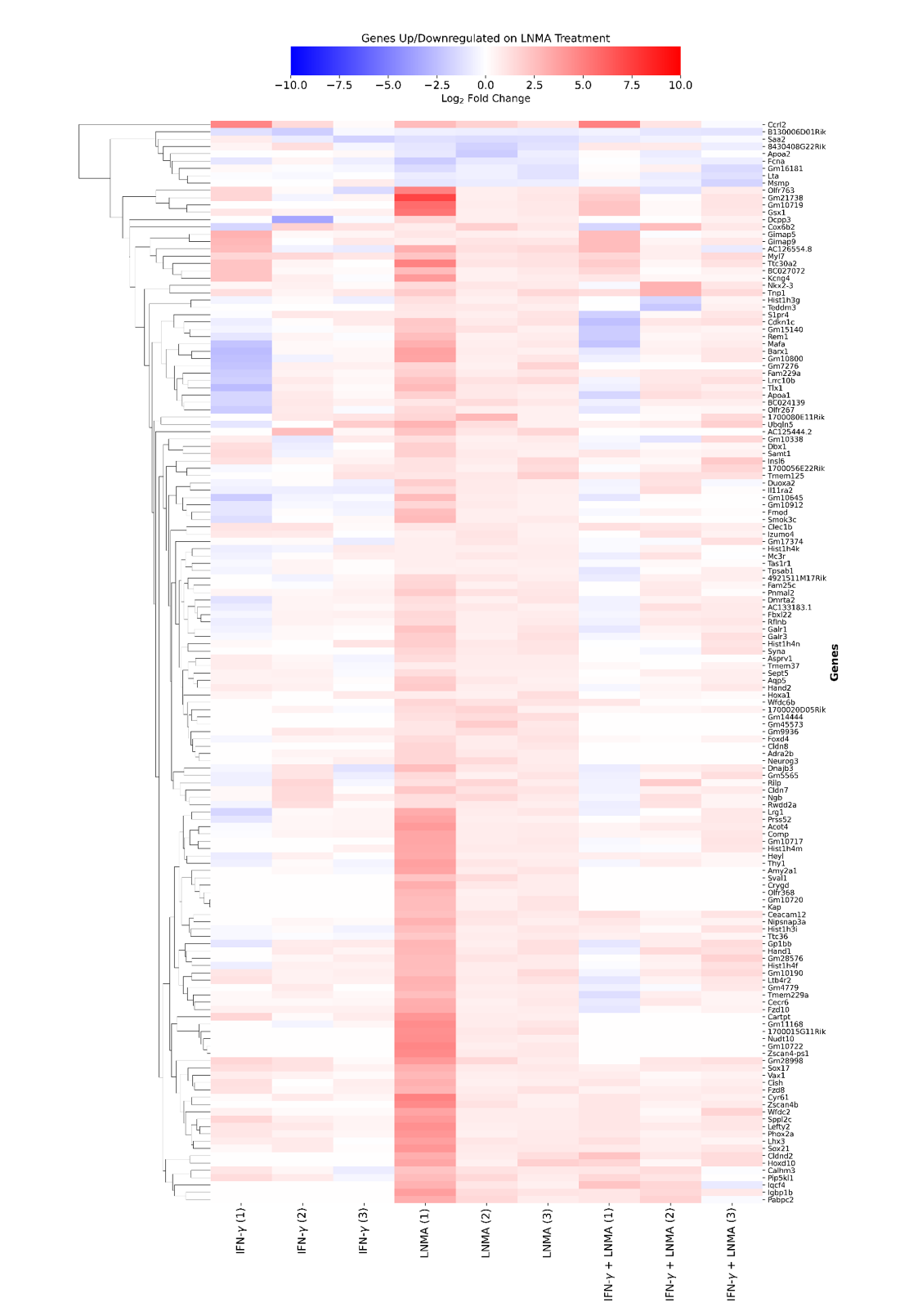

**Fig. S3. Heatmap of DEGs upon LNMA treatment with respect to IFN-γ alone and IFN-γ+LNMA**

Heatmap displaying the log2(fold change) values with respect to control for the LNMA-induced DEGs. The hierarchical clustering was performed based on LNMA-induced DEGs. Data is representative of three independent experiments.

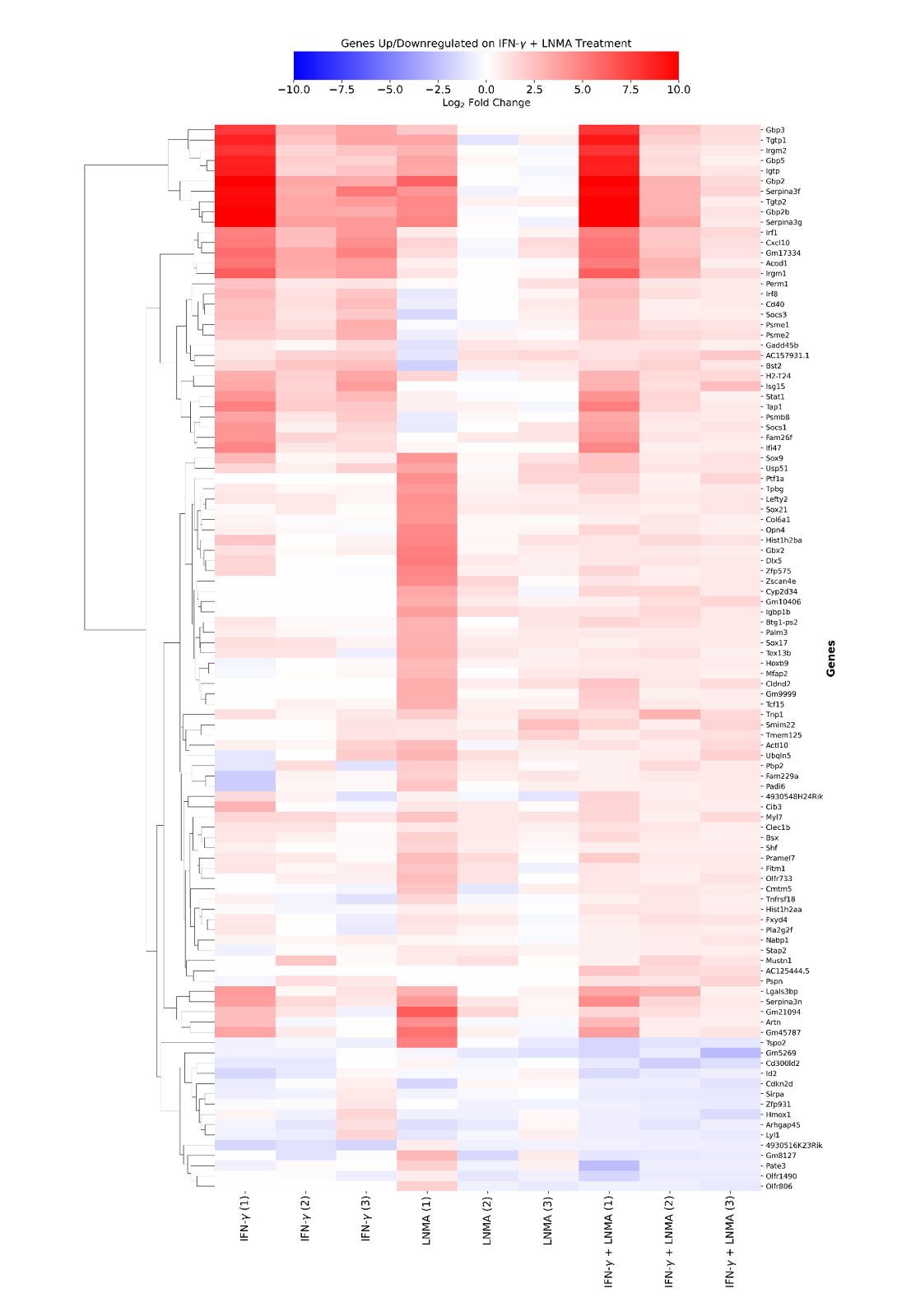

**Fig. S4. Heatmap of DEGs upon IFN-γ+LNMA treatment with respect to treatments of IFN-γ and LNMA alone**

Heatmap displaying the log2(fold change) values with respect to control for the IFN-γ and LNMA-induced DEGs. The hierarchical clustering was performed based on IFN-γ and LNMA-induced DEGs. Data is representative of three independent experiments.

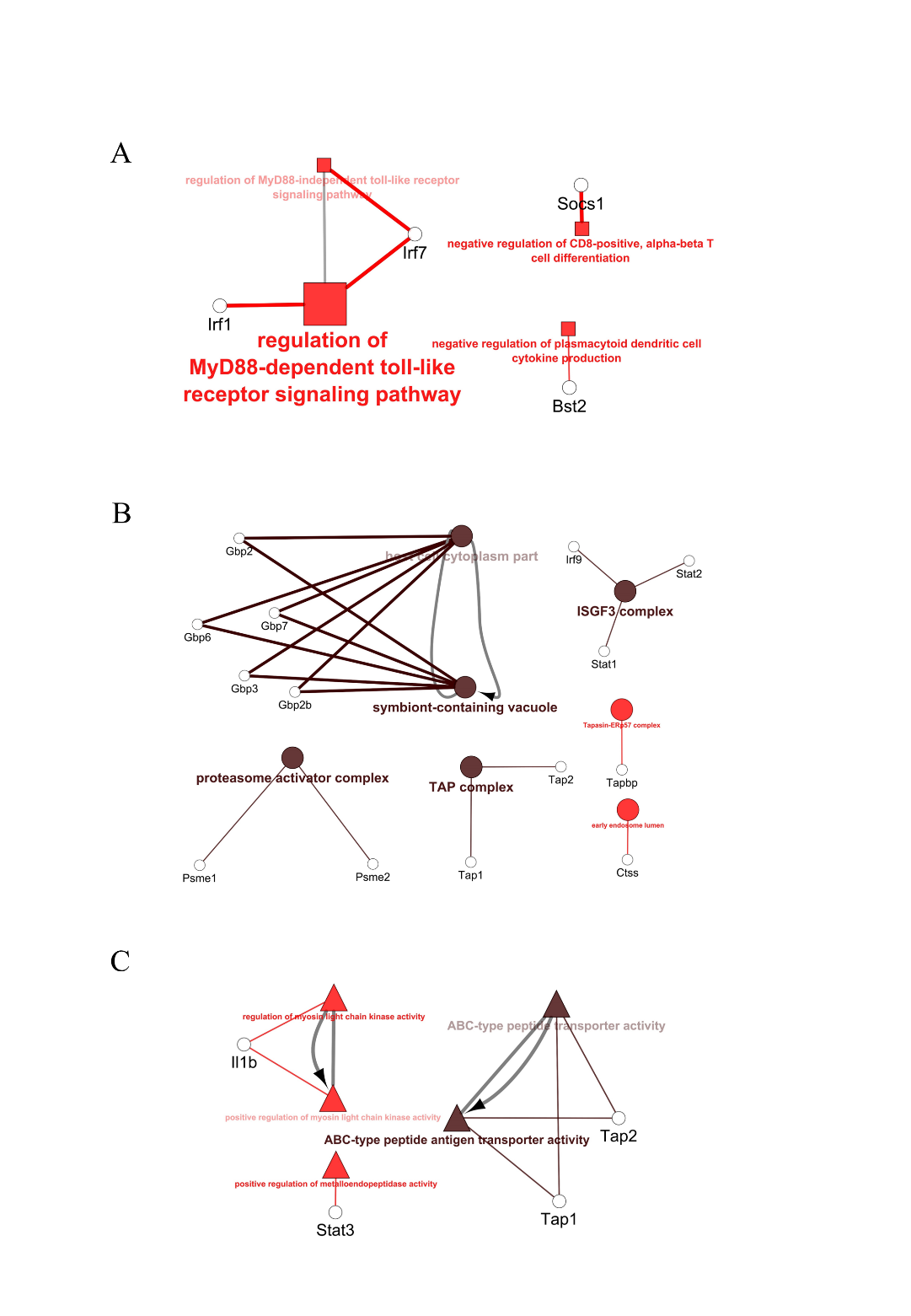

**Fig. S5. Functional enrichment of the toll-like receptor, symbiont-containing vacuole and antigen processing and presentation pathways upon IFN-γ activation**

IFN-γ-induced upregulated DEGs were used as input in the ClueGO plugin of Cytoscape. (A) Immune system, (B) cellular components, and (C) molecular functions of Gene Ontology are displayed. Brown and red color of the nodes represent the Benjamini-Hochberg False Discovery Rate ≤ 0.0001 and ≤0.05, respectively.

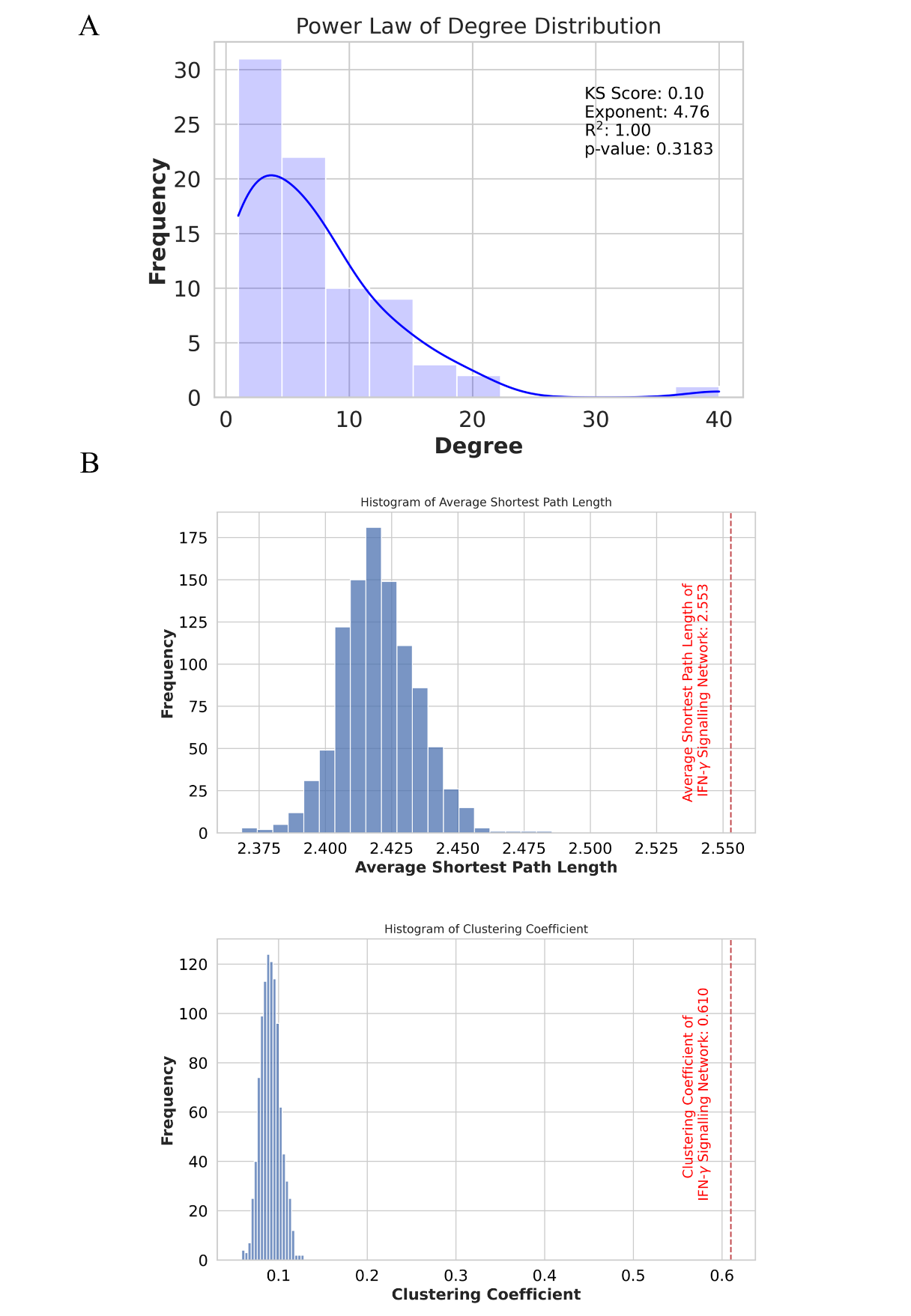

**Fig. S6. IFN-γ-inducible PPI network shows small-world and scale-free properties**

(A) Distribution of node degrees on the Power Law fit analysis. (B) Average shortest path length and clustering coefficient between random networks and the IFN-γ-induced PPI network (red dotted line).

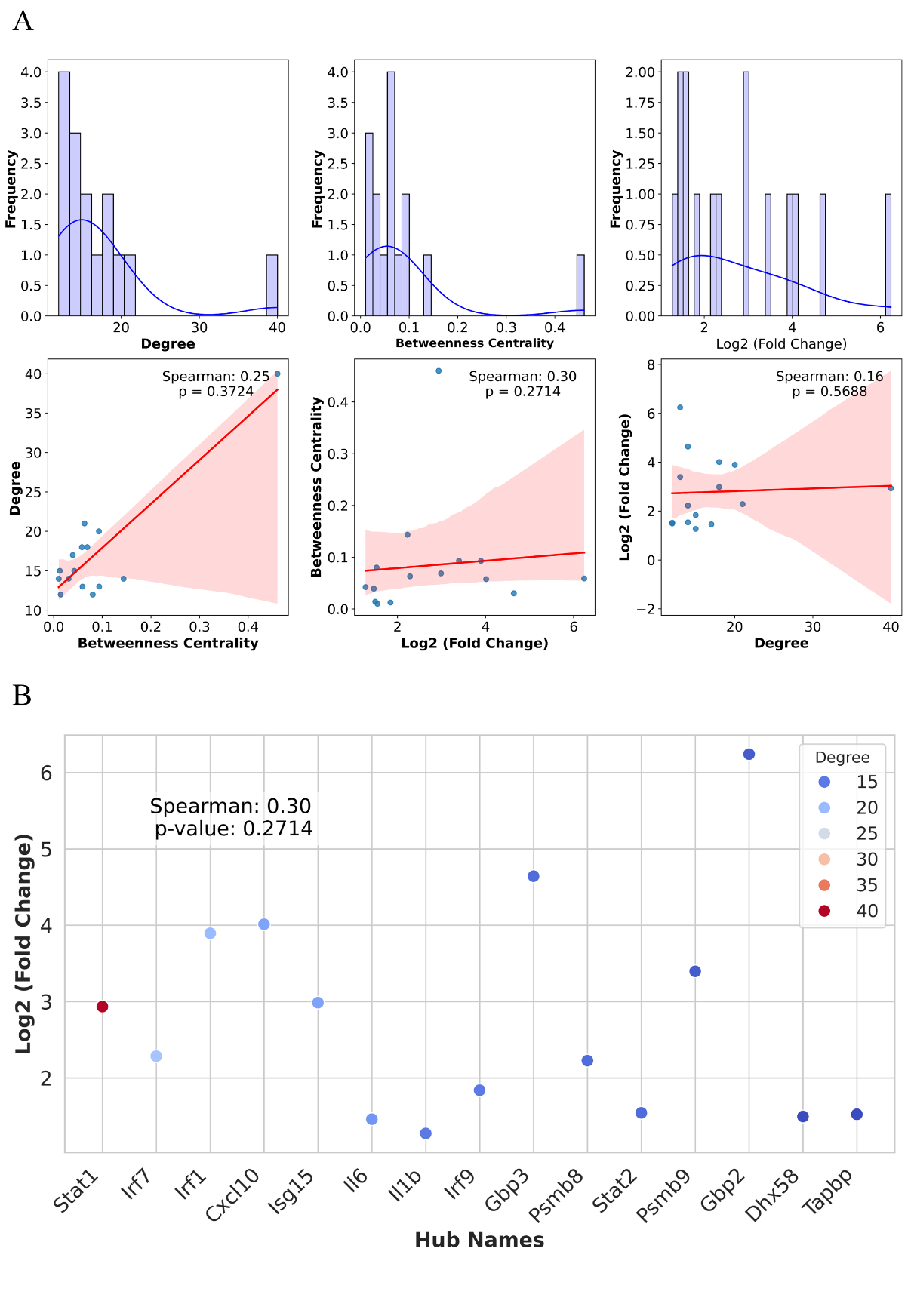

**Fig. S7. Degrees and betweenness centralities of IFN-γ signaling PPI network hubs show no correlation with expression levels but are significantly correlated with each other.**

The KDE plot demonstrates the relationship between degree, betweenness centrality of the PPI network and log2 (fold change) values of the hubs from the PPI network and RNA-seq data.

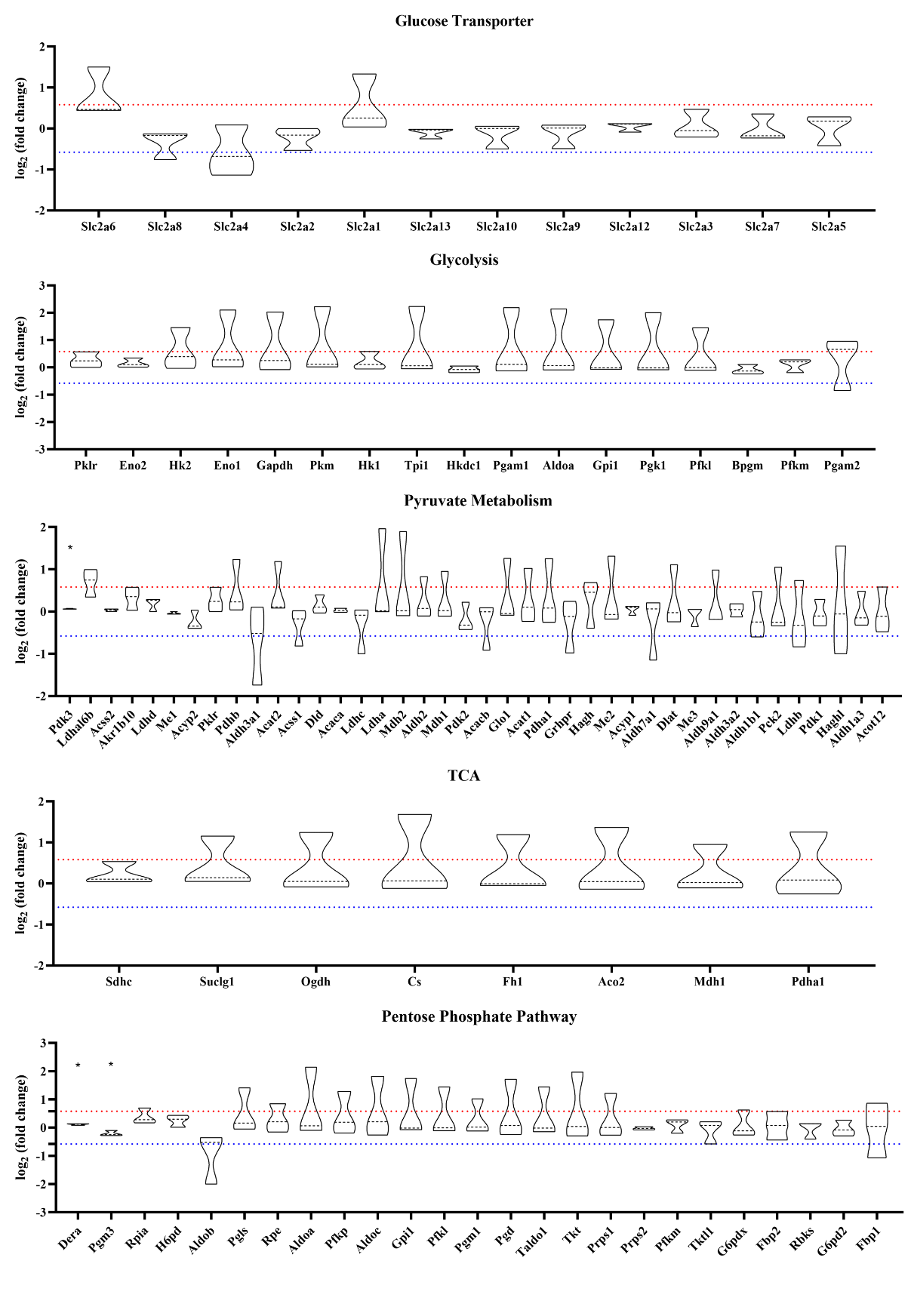

**Fig. S8. Among glucose metabolism genes, glycolytic genes tend to exhibit upregulated expression upon IFN-γ activation.**

'*' indicates a p-value < 0.05, determined by statistical comparison of log2(fold change) values using a T-statistic on Microsoft Excel (version 2404) across three independent experiments.

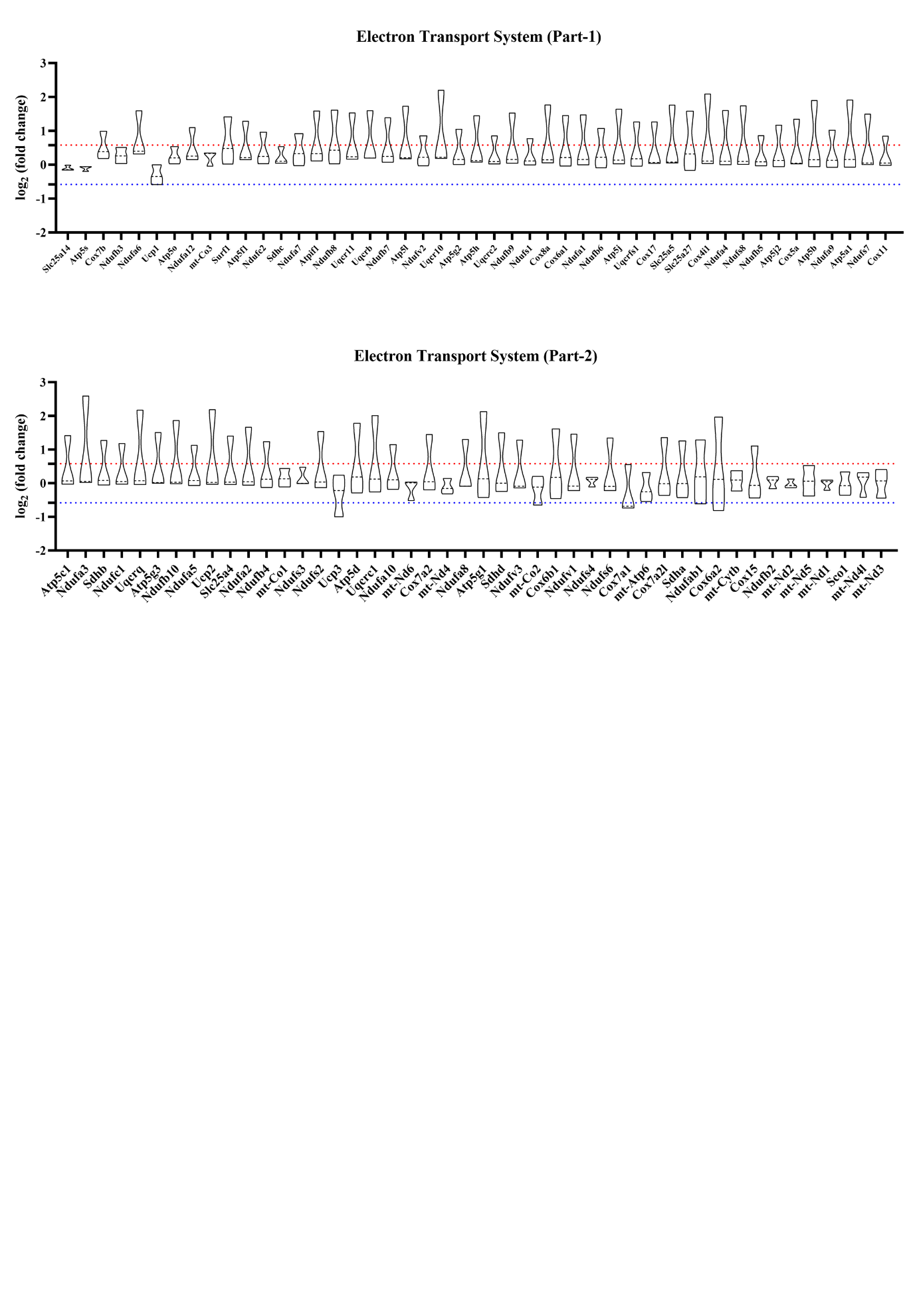

**Fig. S9. Expression of electron transport system genes remain unaffected upon IFN-γ activation**

'*' indicates a p-value < 0.05, determined by statistical comparison of log2(fold change) values using a T-statistic on Microsoft Excel (version 2404) across three independent experiments.

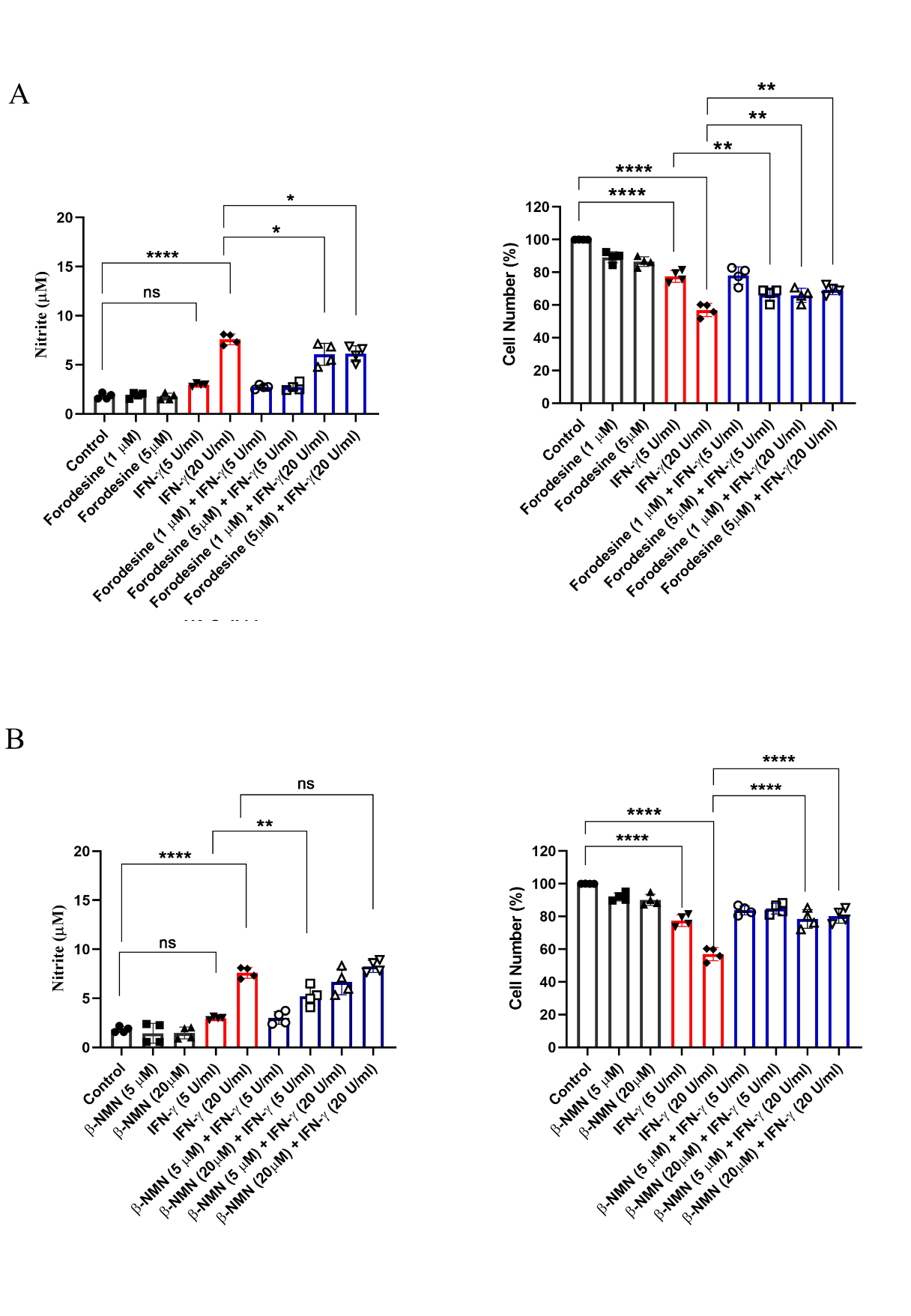

**Fig.S10**. Forodesine (A) was used to inhibit PNP, while β-NMN supplementation (B) served as the NAD^+^ precursor to increase intracellular NAD^+^ concentration. The concentration of nitrite in cell-free supernatant was estimated using Griess assay. The percentage of cell number was measured using Trypan blue dye exclusion assay. The data were acquired after 24 hours of activation. Statistical analyses were performed using ordinary one-way ANOVA with Sidak's multiple comparisons tests. (*), (**), and (****) indicate the statistical differences of p < 0.05, p < 0.01, and p < 0.0001 between the compared groups. Each data point is representative of an independent experiment. Data are represented as mean ± SD of 3-5 independent experiments.

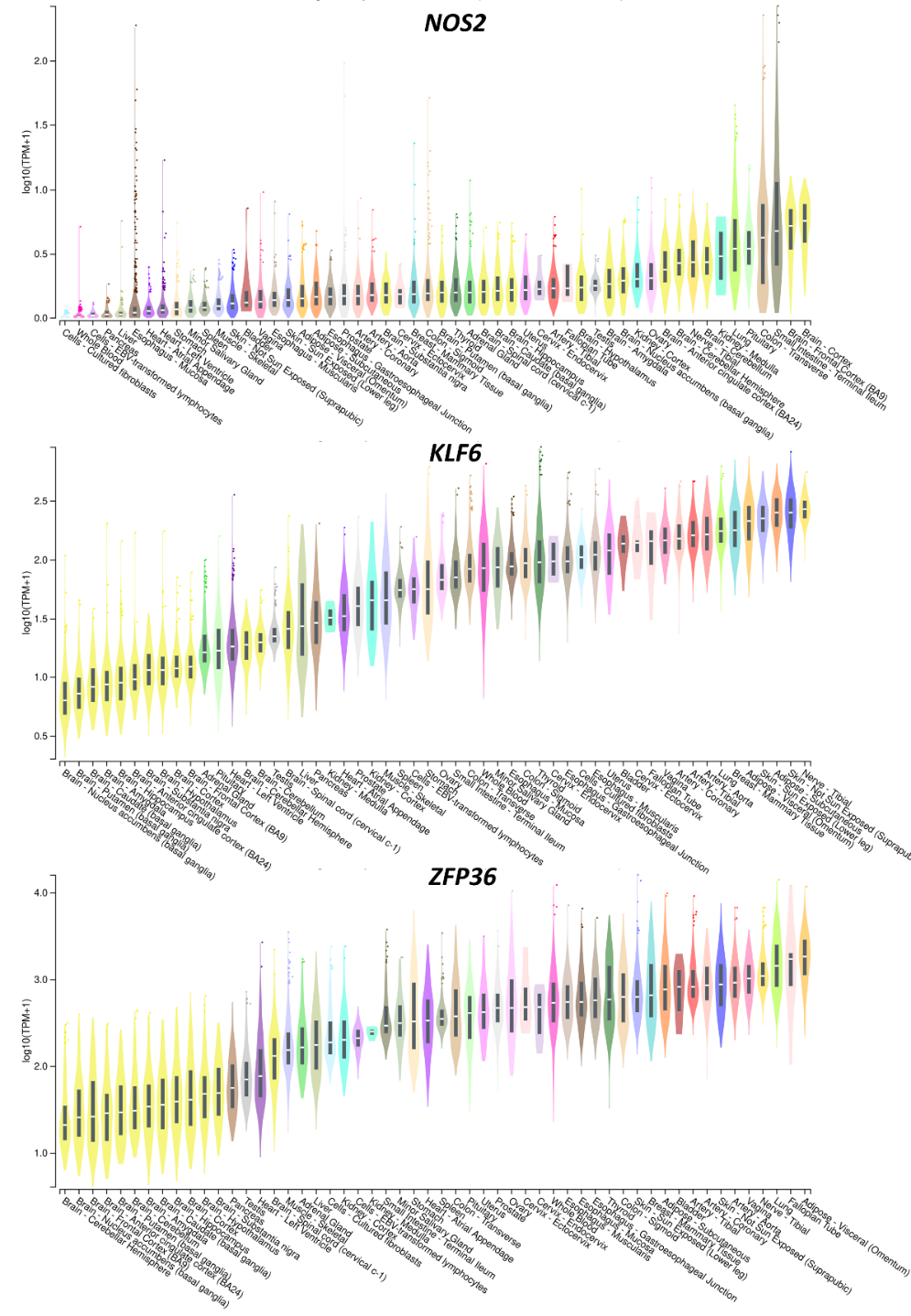

**Fig. S11. Bulk tissue gene expression profile of *NOS2* and NOS-dependent genes: *KLF6* and *ZFP36***

The data was obtained from GTEx Analysis Release V8. Expression values are shown in transcripts per million (TPM), calculated from a gene model with isoforms collapsed to a single gene. No other normalization steps have been applied. Box plots are shown as median and 25^th^ and 75^th^ percentiles; points are displayed as outliers if they are above or below 1.5 times the interquartile range.

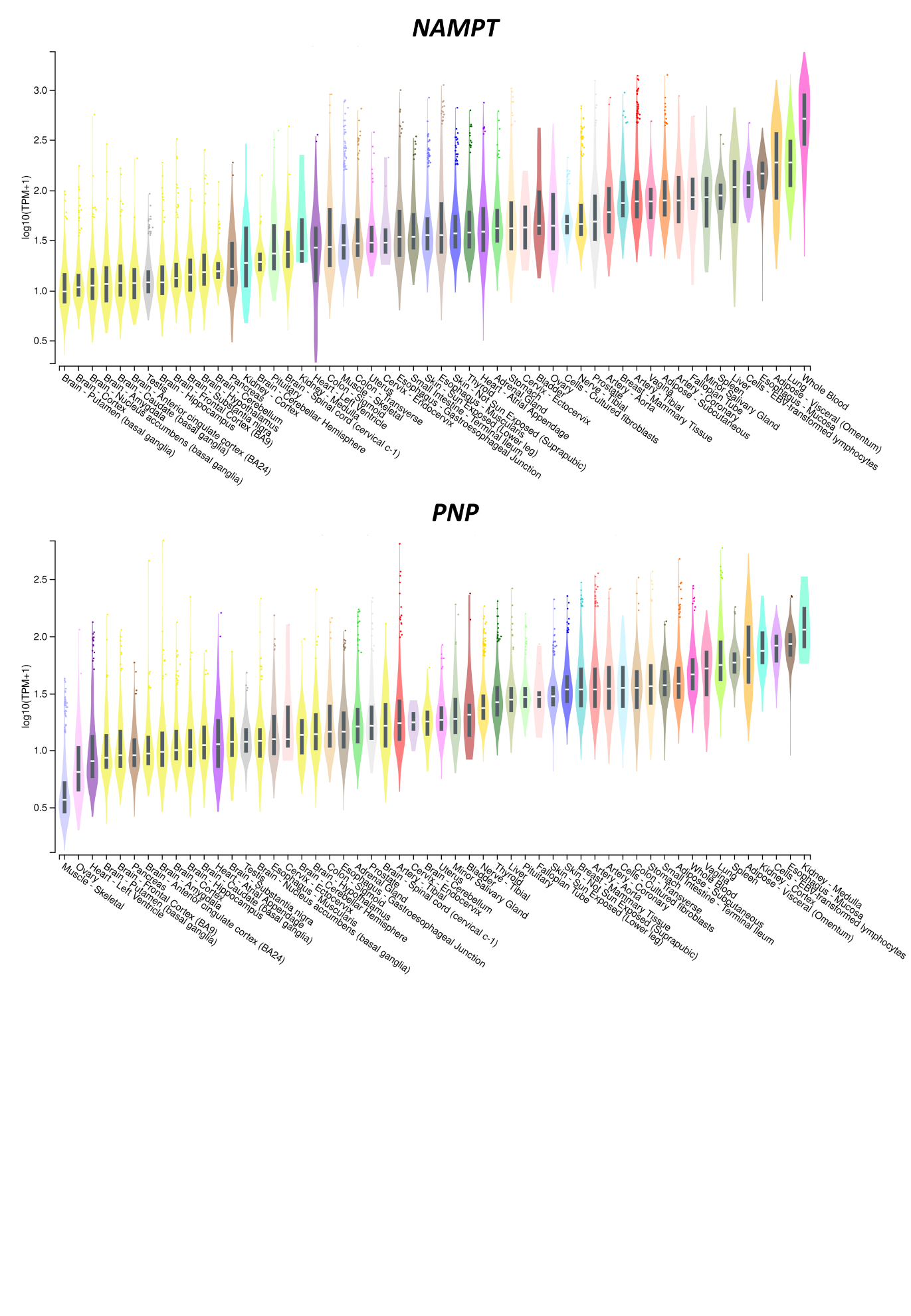

**Fig. S12. Bulk tissue gene expression profile of the nicotinamide metabolism genes: *NAMPT* and *PNP***

The data was obtained from GTEx Analysis Release V8. Expression values are shown in transcripts per million (TPM), calculated from a gene model with isoforms collapsed to a single gene. No other normalization steps have been applied. Box plots are shown as median and 25^th^ and 75^th^ percentiles; points are displayed as outliers if they are above or below 1.5 times the interquartile range.

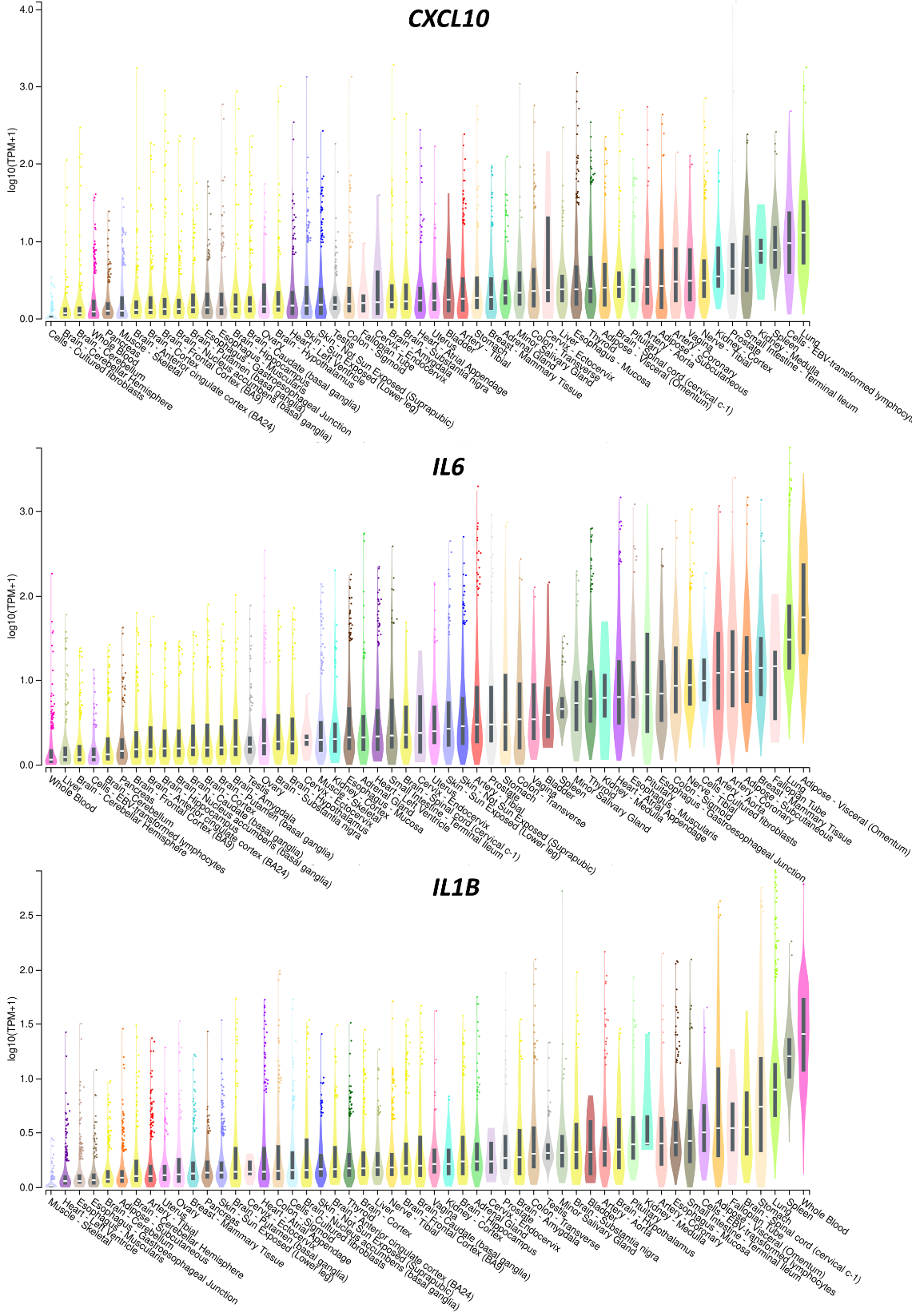

**Fig. S13. The lung is among the top three tissues exhibiting concurrent expression of the cytokine and chemokine hubs *CXCL10*, *IL6*, and *IL1B* under healthy conditions.**

The data was obtained from GTEx Analysis Release V8. Expression values are shown in transcripts per million (TPM), calculated from a gene model with isoforms collapsed to a single gene. No other normalization steps have been applied. Box plots are shown as median and 25^th^ and 75^th^ percentiles; points are displayed as outliers if they are above or below 1.5 times the interquartile range.

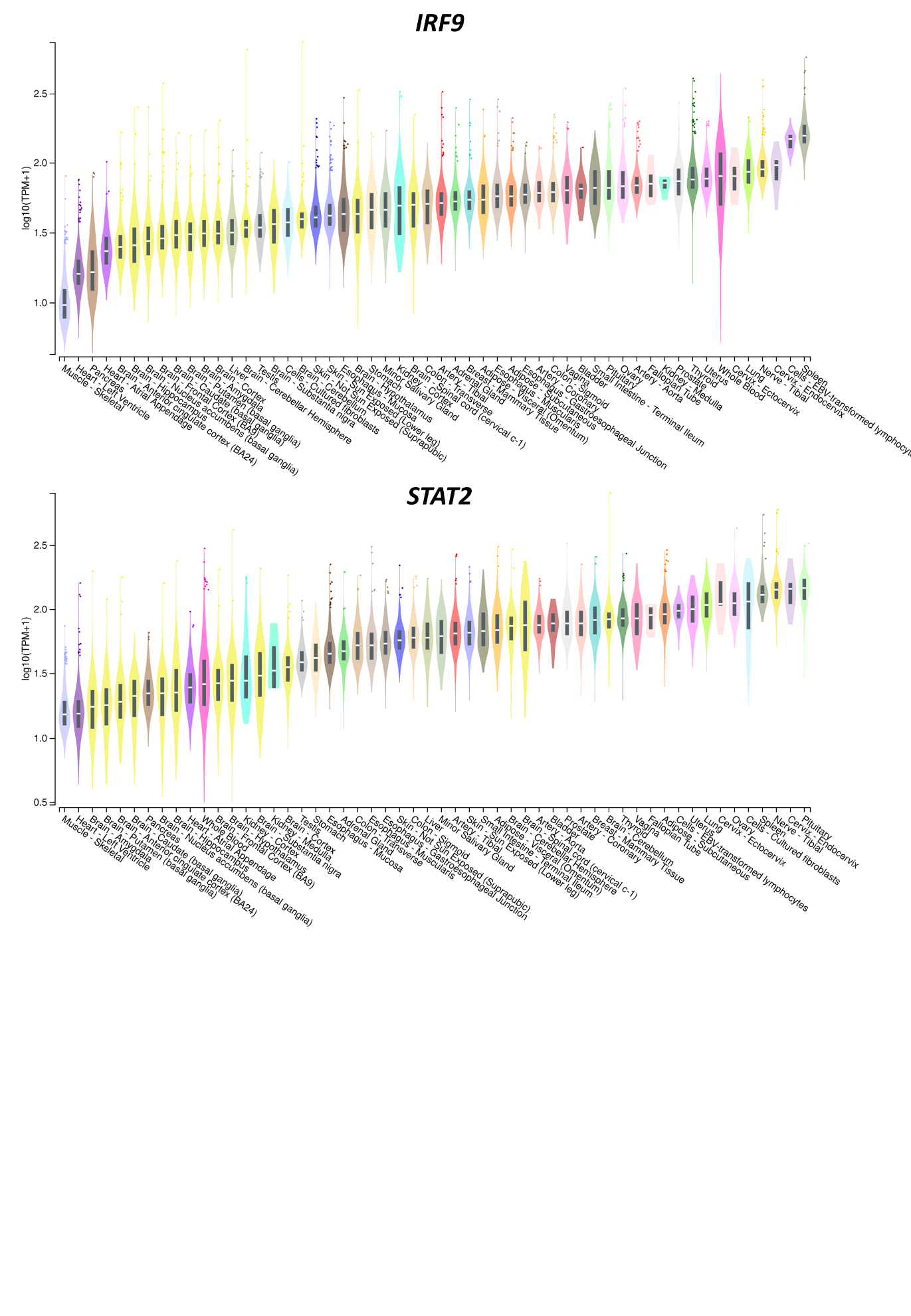

**Fig. S14. Bulk tissue gene expression profile of the hubs: *IRF9* and *STAT2***

The data was obtained from GTEx Analysis Release V8. Expression values are shown in transcripts per million (TPM), calculated from a gene model with isoforms collapsed to a single gene. No other normalization steps have been applied. Box plots are shown as median and 25^th^ and 75^th^ percentiles; points are displayed as outliers if they are above or below 1.5 times the interquartile range.

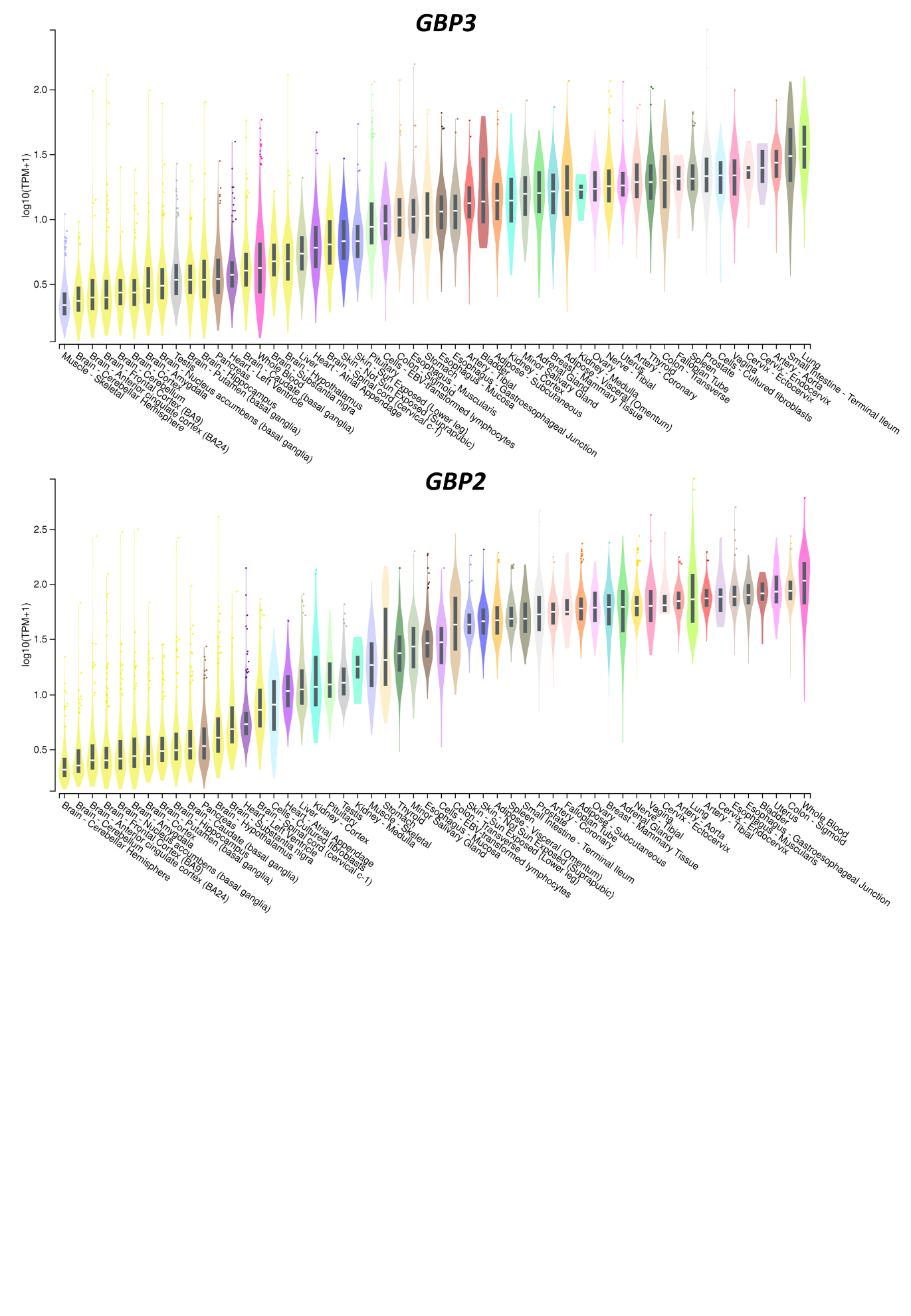

**Fig. S15. Bulk tissue gene expression profile of the GBP hubs: *GBP3 and GBP2***

The data was obtained from GTEx Analysis Release V8. Expression values are shown in transcripts per million (TPM), calculated from a gene model with isoforms collapsed to a single gene. No other normalization steps have been applied. Box plots are shown as median and 25^th^ and 75^th^ percentiles; points are displayed as outliers if they are above or below 1.5 times the interquartile range.

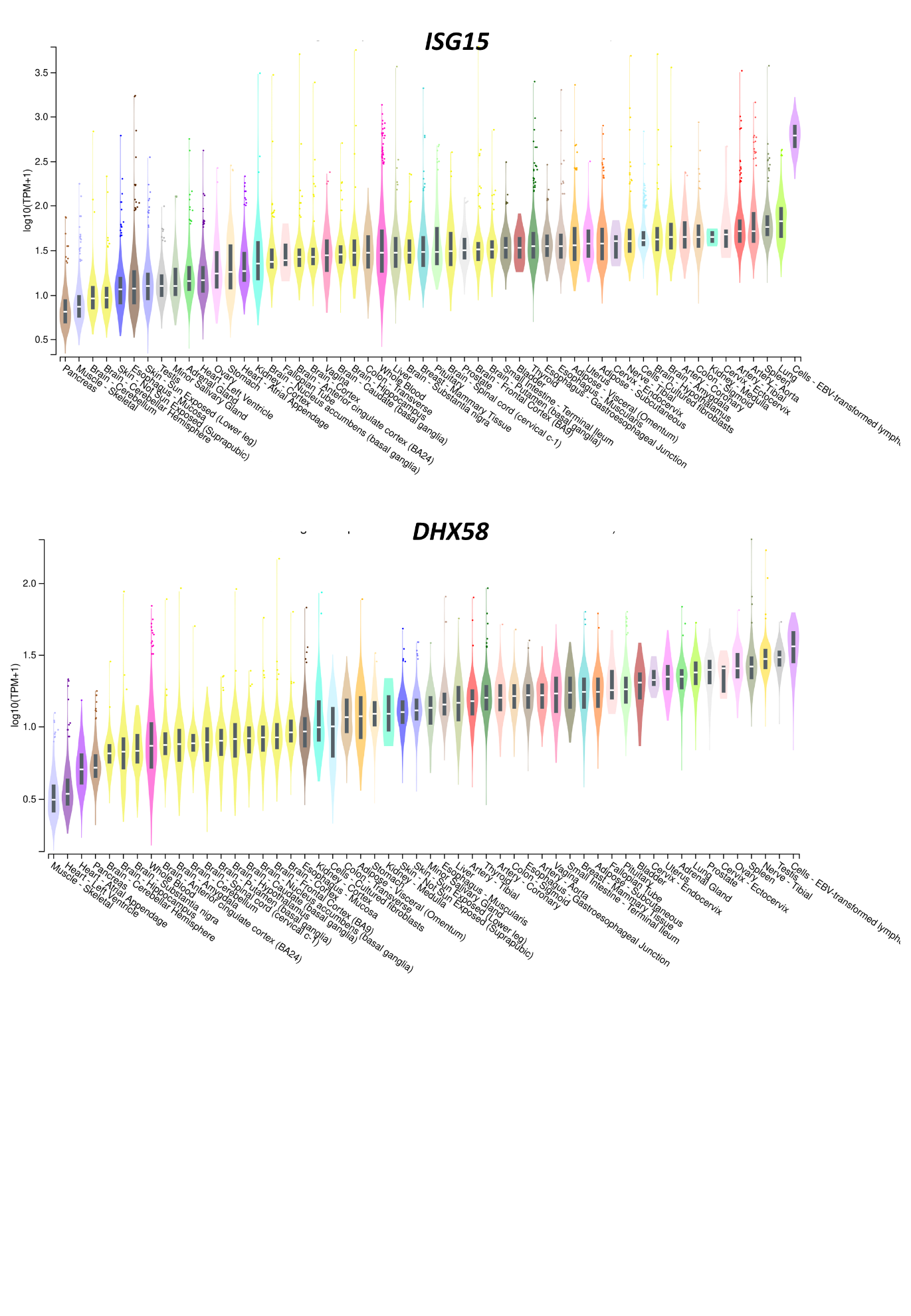

**Fig. S16. EBV-transformed lymphocytes topped in expressing the hubs: *ISG15* and *DHX58***

The data was obtained from GTEx Analysis Release V8. Expression values are shown in transcripts per million (TPM), calculated from a gene model with isoforms collapsed to a single gene. No other normalization steps have been applied. Box plots are shown as median and 25^th^ and 75^th^ percentiles; points are displayed as outliers if they are above or below 1.5 times the interquartile range.

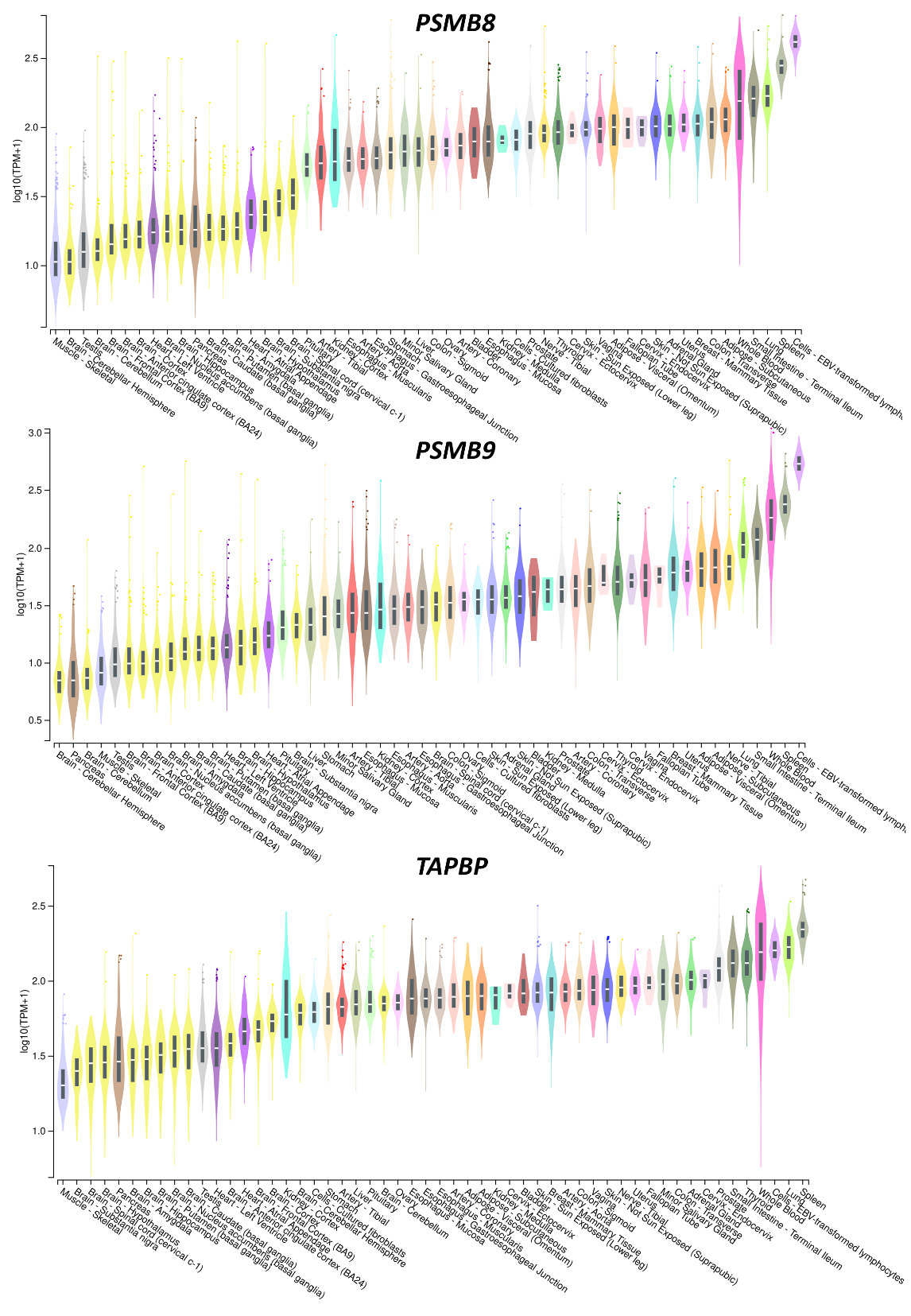

**Fig. S17. EBV-transformed lymphocytes, lung, and spleen exhibited the highest expression of the antigen processing and presentation hubs *PSMB8*, *PSMB9*, and *TAPBP***

The data was obtained from GTEx Analysis Release V8. Expression values are shown in transcripts per million (TPM), calculated from a gene model with isoforms collapsed to a single gene. No other normalization steps have been applied. Box plots are shown as median and 25^th^ and 75^th^ percentiles; points are displayed as outliers if they are above or below 1.5 times the interquartile range.
